# Using a Quadruplet Codon to Expand the Genetic Code of an Animal

**DOI:** 10.1101/2021.07.17.452788

**Authors:** Zhiyan Xi, Lloyd Davis, Kieran Baxter, Ailish Tynan, Angeliki Goutou, Sebastian Greiss

## Abstract

Genetic code expansion in multicellular organisms is currently limited to the use of repurposed amber stop codons. Here we introduce a system for the use of quadruplet codons to direct incorporation of non-canonical amino acids *in vivo* in an animal, the nematode worm *Caenorhabditis elegans*. We develop hybrid pyrrolysyl tRNA variants to incorporate non-canonical amino acids in response to the quadruplet codon UAGA. We demonstrate the efficiency of the quadruplet decoding system by incorporating photocaged amino acids into two proteins widely used as genetic tools. We use photocaged lysine to express photocaged Cre recombinase for the optical control of gene expression and photocaged cysteine to express photo-activatable caspase for light inducible cell ablation. Our approach will facilitate the routine adoption of quadruplet decoding for genetic code expansion in eukaryotic cells and multicellular organisms.

## Introduction

The genetic code, universally shared among all kingdoms of life, is based on nucleotide triplet codons. Of the 64 possible triplet permutations, 61 are sense codons that map to 20 canonical amino acids and 3 are stop codons to terminate translation [1]. Expanding the genetic code beyond the 20 canonical amino acids requires the reassignment of at least one codon to code for an additional amino acid. Examples in nature are the assignments of the stop codon UGA to encode selenocysteine, which is found in all three kingdoms of life, or UAG to encode pyrrolysine in methanogenic archaea and bacteria [2–5].

The development of genetic code expansion technology for the artificial expansion of the genetic code has allowed the incorporation of a wide range of synthetic non-canonical amino acids (ncAA) with properties not found in nature [6]. The method involves the addition of an orthogonal aminoacyl-tRNA-synthetase (aaRS) / tRNA pair to the translational machinery of a cell. The aaRS specifically recognises the ncAA and attaches it to the tRNA. At the ribosome the charged tRNA then pairs with a matching codon, inserted in the reading frame of a target gene to determine the site of incorporation. Genetic code expansion has been established in a number of organisms, including single celled systems such as bacteria, yeast, mammalian cell culture, as well as multicellular organisms including *C. elegans*, *Drosophila*, zebrafish and mouse [7–12]. The codon most widely used to direct ncAA incorporation is the amber stop codon UAG, and in multicellular organisms UAG is the only codon that has thus far been used for this purpose.

Genetic code expansion depends on the availability of coding space in the form of nucleotide codons to direct incorporation, with one codon required for each ncAA to be used independently in a cell. Strategies to expand coding space include genomic recoding to free up and then reassign sense codons, which is being explored in prokaryotic cells [13, 14]. However, this approach is not only impractical in more complex systems, but also undesirable, as extensive genomic recoding may disrupt the normal biological function of a model system used to study biological processes. Another approach, the use of quadruplet codons does not require recoding and offers a potential expansion from 64 to 256 codons. The use of quadruplet codons has thus far however only been possible in single celled systems [15, 16]. The development of an efficient quadruplet decoding system in multicellular organisms would help to open a path towards further expanding the utility of ncAA mutagenesis, especially in complex systems. Apart from the prospect of allowing the concurrent use of multiple ncAA in the same cell or even the same protein, a further benefit of quadruplet codons may lie in the reduction of potential cross-decoding of endogenous triplet codons. The use of the triplet tRNA_CUA_, that recognises the UAG stop codon, has been shown to result in incorporation of ncAA at endogenous UAG stop codons, an effect that can be reduced when employing the UAGA quadruplet codon [17, 18].

Despite the advantages offered by quadruplet codons, their use has been held back due to the low ncAA incorporation efficiency compared with triplet stop codons [17, 19, 20]. Attempts to improve quadruplet decoding have focused on the directed evolution of anticodon loops for use in bacteria and mammalian cell culture [16, 18, 21], and on the evolution of the ribosomal decoding centre in *E. coli* [22].

Here we present the first method for quadruplet decoding in a multicellular organism, the nematode worm *C. elegans*. We develop hybrid tRNA variants by fusing anticodon loops optimised for quadruplet decoding [16, 18, 23] to tRNA scaffolds optimised for interaction with the eukaryotic translational machinery [24, 25]. We use our system to incorporate two distinct ncAAs, photocaged lysine (PCK) and photocaged cysteine (PCC), and show that we can achieve incorporation efficiencies that come close to those achieved using UAG triplet codons. We demonstrate the utility of the system by using quadruplet codons i) to incorporate PCK site specifically into Cre recombinase to express a photo-activatable version of Cre which we use to optically control gene expression in live animals, and ii) to incorporate PCC into a constitutively active version of Caspase-3 to engineer a photo-activatable caspase, which we use to optically ablate *C. elegans* neurons.

This advance demonstrates the feasibility of going beyond the standard triplet based genetic code in complex multicellular systems, and paves the way for harnessing the advantages of quadruplet decoding for genetic code expansion applications in multicellular organisms.

## Results

### Efficient quadruplet decoding in *C. elegans*

Quadruplet decoding requires tRNA molecules that possess the ability to introduce +1 frameshifts during decoding, yet can function efficiently within the endogenous translational machinery which is evolutionarily optimised towards triplet codons and against translational frameshifts.

Modifications introduced into the tRNA anticodon loop outside of the anticodon can significantly improve the capability of tRNAs to introduce a +1 frameshift when delivering an amino acid to the ribosome [18]. Directed evolution approaches have been used to generate improved anticodon loops, which have been used for quadruplet decoding in bacteria and cultured mammalian cells [16, 18, 21].

We aimed to construct a system for quadruplet decoding in *C. elegans* based on the pyrrolysyl-tRNA synthetase (PylRS) / tRNA(Pyl) pair from *Methanosarcina* species, which is functional in *C. elegans* [10, 25] and has been used for quadruplet decoding in both bacteria and cultured mammalian cells [16, 18, 21].

To allow comparison of incorporation efficiency between a quadruplet decoding system and the established *C. elegans* UAG triplet decoding system, we decided to use the UAGA quadruplet codon, which has been used for ncAA incorporation in eukaryotic cells [23] and shares its first three nucleotides with UAG (Figure 1).

**Figure 1.**
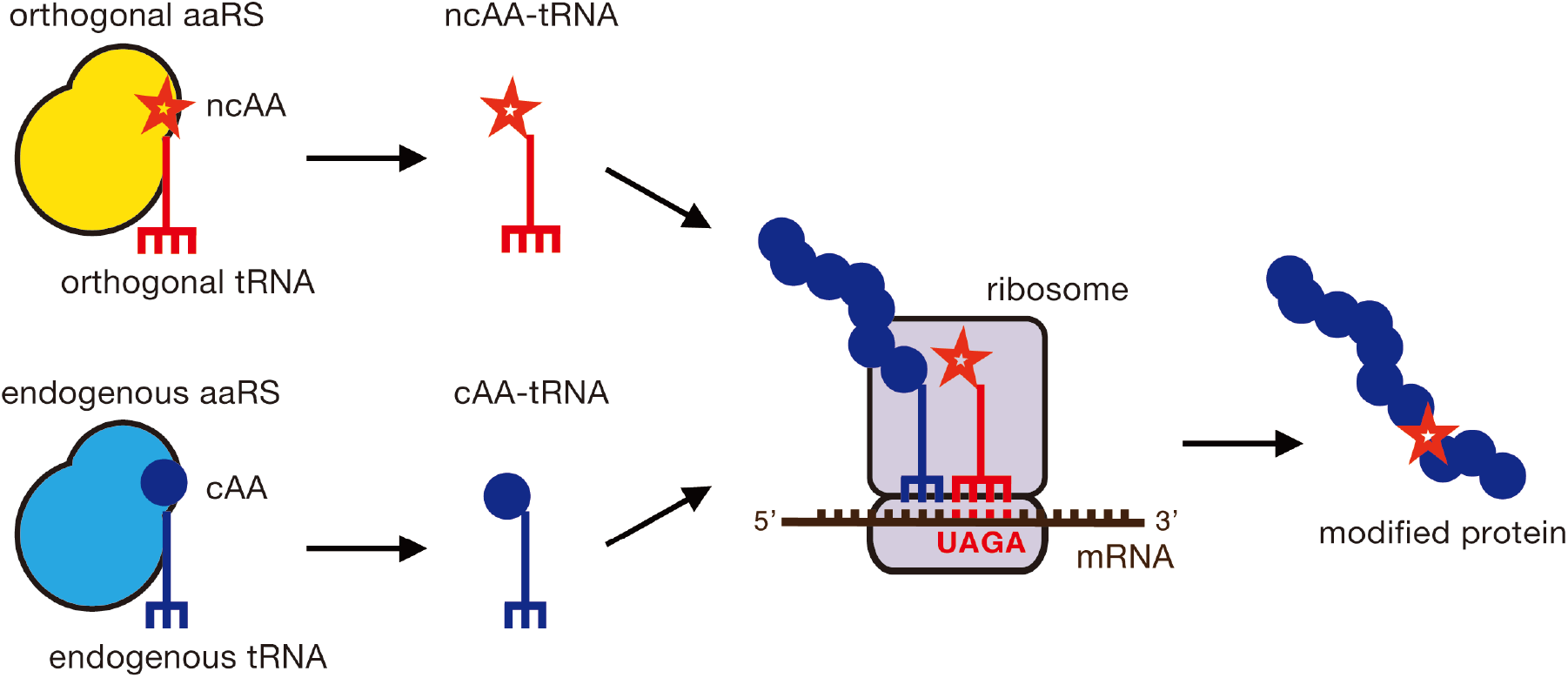
Quadruplet decoding for the incorporation of non-canonical amino acids. An orthogonal aminoacyl-tRNA synthetase (aaRS) attaches a non-canonical amino acid (ncAA) to its cognate tRNA. The tRNA contains a quadruplet anticodon. The orthogonal components do not interact with endogenous amino acids, aaRSs or tRNAs. At the ribosome, the ncAA is incorporated into the growing polypeptide chain in response to a quadruplet codon during translation.

We first tested two tRNA(Pyl) variants, tRNA(Pyl/M7) and tRNA(Pyl/UAGA-1) which possess anticodon loops optimised for quadruplet decoding. The tRNA(Pyl/M7) contains the M7 anticodon loop and was isolated from a pool of tRNA(Pyl) variants carrying the UCCU anticodon that decodes AGGA [16], while the UAGA-1 anticodon loop was isolated from a pool of tRNA(Pyl) variants in a screen for improved UAGA decoding [18, 23]. Since the anticodon loop of tRNA(Pyl/M7) differs from tRNA(Pyl/UAGA-1) in only 3 nucleotides outside the anticodon and 3 out of its 6 differences compared to the wild type tRNA(Pyl)_CUA_ sequence are conserved between M7 and UAGA-1, we surmised that the M7 variant might be effective even when the anticodon is changed from UCCU to UCUA for decoding UAGA (Supplementary Table1).

To test the functionality of the quadruplet decoding tRNAs, we constructed transgenic *C. elegans* lines carrying three plasmids (Figure 2A): i) a plasmid expressing the PylRS variant NES::PCKRS that recognises photocaged lysine (PCK) [25, 26] and at its N-terminus contains a nuclear export signal (NES) derived from human Smad4 [27], which dramatically improves the efficiency of the enzyme [25, 28]; ii) tRNA(Pyl/M7)_UCUA_ or tRNA(Pyl/UAGA-1)_UCUA_ with expression driven by the ubiquitous RNA polymerase III promoter *rpr-1p* [29]; and iii) a fluorescent reporter for PCK incorporation. The fluorescent reporter consists of a GFP gene separated by a UAGA codon from a downstream mCherry gene, followed by an HA tag and a nuclear localisation sequence (NLS) (Figure 2A). Incorporation at the UAGA codon thus results in production of a GFP::mCherry fusion protein that localises to the cell nucleus due to the C-terminal NLS. Expression of the protein coding genes was driven by the ubiquitous *sur-5p* promoter [30]. To construct strains, we used a genetic background containing a deletion in the *smg-6* gene, knocking out the nonsense mediated decay machinery. This dysfunction in nonsense mediated decay helps to stabilise the reporter mRNA. All transgenic strains were generated by biolistic bombardment [31].

**Figure 2.**
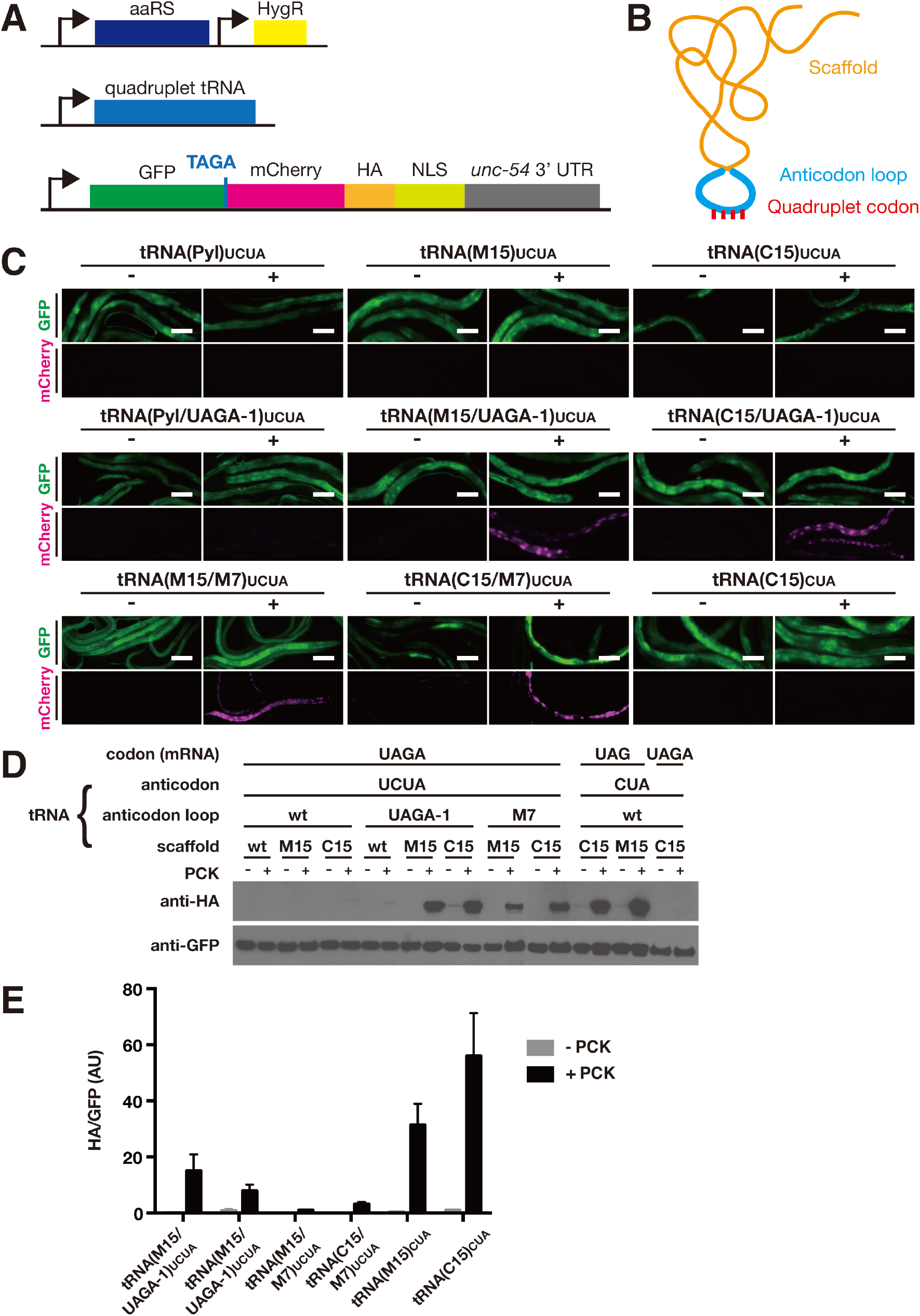
Efficient quadruplet decoding using hybrid tRNAs. **(A)** Genetic constructs for the quadruplet decoding machinery consisting of aaRS and quadruplet tRNA. The machinery is co-expressed with a fluorescent reporter for ncAA incorporation. **(B)** Schematic of hybrid tRNA consisting of scaffold and anticodon loop. **(C)** Fluorescence microscopy images of randomly selected worms expressing different tRNA variants together with NES::PCKRS, grown in the presence (“+”) or absence (“−”) of 1 mM PCK. “Pyl”, “M15” and “C15” denote scaffolds, “UAGA-1” and “M7” denote anticodon loops. Scale bar = 100 μm. **(D)** Western blots of lysates made from transgenic lines shown in (C). Full-length reporter protein was detected using anti-HA antibody, samples were normalised using anti-GFP antibody. **(E)** Quantitative western blots of the constructs showing the highest levels of incorporation in (C) and (D). For all constructs, two independent lines were quantified.

To assess PCK incorporation, we grew transgenic animals for 48h on nematode growth medium (NGM) agar plates supplemented with 1 mM PCK. We observed very weak, barely visible mCherry fluorescence indicative of PCK incorporation in a minority of animals when using tRNA(Pyl/UAGA-1)_UCUA_, (Figure 2C). We confirmed presence of the full-length GFP::mCherry::HA product following incorporation by western blot using an anti-HA antibody to detect the C-terminal HA tag, albeit a band was only visible after extended exposure of the blot to film (Figure 2D & Supplementary Figure 1). As expected, no incorporation was detected in the absence of PCK. In contrast to tRNA(Pyl/UAGA-1)_UCUA_, we observed no incorporation when using tRNA(Pyl/M7)_UCUA_, or tRNA(Pyl)_UCUA_, which contains the UCUA anticodon in a wild type anticodon loop (Figure 2C,D & Supplementary Figure 1).

### Optimised hybrid tRNAs for quadruplet decoding

The archeal tRNA(Pyl) contains sequence elements that differ from canonical mammalian tRNAs and these structural differences disfavour the interaction of tRNA(Pyl) with the eukaryotic translational machinery [24]. The efficiency of ncAA incorporation in mammalian cells and in *C. elegans* can be improved many-fold by the introduction of facilitative structural elements into the tRNA scaffold [24, 25]. As these optimising tRNA mutations are outside the anticodon loop, we set out to assess the possibility of constructing hybrid tRNAs for quadruplet decoding by combining optimised anticodon loops with scaffolds optimised for the eukaryotic translational context (Figure 2B).

We based the new hybrid tRNAs on two optimised scaffolds originally engineered for usage in mammalian cells, M15 and C15, which drastically increase ncAA incorporation efficiency at UAG triplet codons in *C. elegans* [25]. We combined the M15 and C15 scaffolds with the M7 and UAGA-1 anticodon loops to create four hybrid tRNA variants: tRNA(M15/M7)_UCUA_, tRNA(C15/M7)_UCUA_, tRNA(M15/UAGA-1)_UCUA_, and tRNA(C15/UAGA-1)_UCUA_. All four hybrid variants contained the UCUA anticodon to decode the UAGA quadruplet codon. We generated transgenic strains co-expressing the hybrid tRNA variants together with the synthetase NES::PCKRS and the GFP::UAGA::mCherry reporter (Figure 2A).

We then tested incorporation efficiencies by growing transgenic animals on NGM plates supplemented with 1 mM PCK. In the strains expressing hybrid tRNAs, we saw a striking improvement in incorporation efficiency as compared to the strains expressing tRNAs with only the anticodon loops optimised (Figure 2C, D). The UAGA-1 anticodon loop appeared to outperform the M7 anticodon loop, while both the M15 and the C15 scaffolds showed comparable improvements to incorporation efficiency. The improvement in incorporation rates was due to the combined effect of optimised scaffold and loops, since tRNA(C15)_UCUA_, which combines an optimised scaffold with the wild type anticodon loop showed only barely detectable incorporation (Figure 2D).

We performed quantitative western blots comparing incorporation rates of the 4 hybrid quadruplet tRNAs against triplet codon incorporation rates achieved using the parent tRNA molecules tRNA(M15)_CUA_ and tRNA(C15)_CUA_ incorporating at a UAG triplet codon. While we observed the highest incorporation rates when using the triplet incorporation system, the best performing quadruplet tRNA(M15/UAGA-1) reached 30-50% of the levels of incorporation observed for the triplet tRNAs M15 and C15 (Figure 2C, D). Strikingly, the best performing quadruplet tRNA(C15/UAGA-1) showed a 4-fold improvement of incorporation compared to the standard, non-optimised UAG triplet system consisting of wild type tRNA(Pyl)_CUA_ and PCKRS without an N-terminal nuclear export sequence (Supplementary Figure 2). Due to the high incorporation efficiency of the improved quadruplet system, we decided to perform all further experiments in the wild type N2 genetic background with functional nonsense mediated decay.

### Application of quadruplet decoding for the expression of photocaged Cre recombinase and optical control of gene expression in *C. elegans*

We next decided to assess the utility of the optimised quadruplet-decoding system by using it to express a light controllable version of Cre recombinase. Lysine residue K201 in the catalytic site of Cre recombinase is required for activity and replacing it with PCK renders the enzyme inactive. Uncaging by short illumination with 365 nm light restores Cre activity and can thus be used to activate Cre and switch on expression of Cre target genes [32] (Figure 3B). We have previously established photocaged Cre as a tool for optical control of gene expression in *C. elegans* [25], using a UAG triplet codon system to direct incorporation of PCK. Here, instead of UAG, we used the UAGA quadruplet codon to direct incorporation of PCK in place of the active site lysine K201. To decode UAGA, we tested the two quadruplet tRNA variants based on the M15 scaffold, tRNA(M15/M7)_UCUA_ and tRNA(M15/UAGA-1)_UCUA_ (Figure 3C).

**Figure 3.**
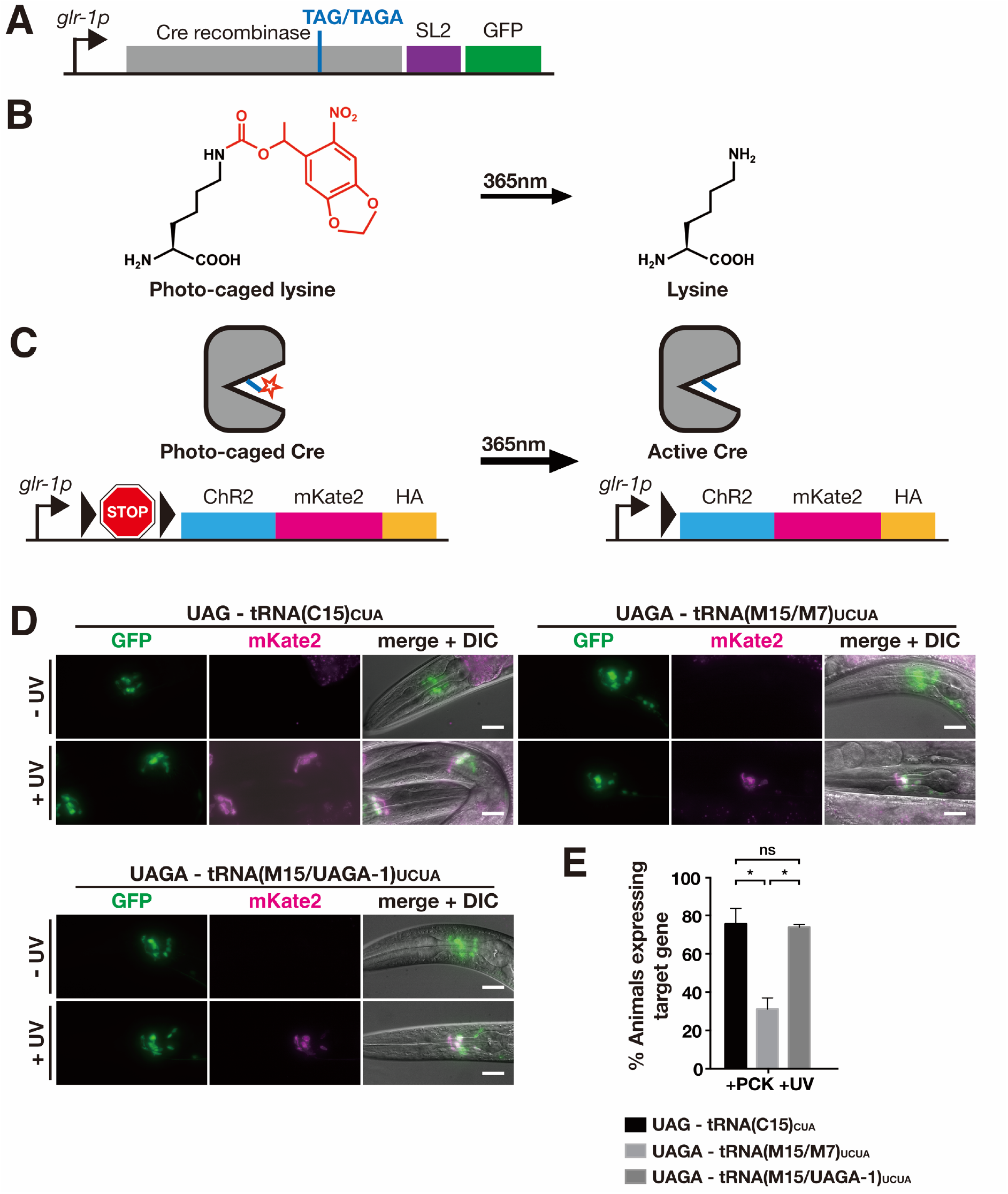
Using quadruplet codons to express photocaged Cre recombinase. **(A)** Genetic constructs for expression of photocaged Cre. Photocaged lysine PCK replaces the active site lysine K201 of Cre. The incorporation site is specified using either a TAG or TAGA codon. GFP is co-expressed with photocaged Cre. **(B)** 6-nitropiperonyl-L-lysine, “photocaged lysine”. The caging group is stable in visible light and can be removed by brief illumination with 365 nm light. **(C)** Optical activation of Cre. When introduced in place of K201, the caging group of PCK (red star) blocks Cre activity. Transcription of the target gene is blocked by a transcriptional terminator (“STOP”) inserted between the gene and its promoter. The terminator is flanked by loxP sites (black triangles). Uncaging activates Cre, which removes the transcriptional terminator, thereby activating target gene expression. **(D)** Fluorescence microscopy images of transgenic lines expressing the PCK incorporation machinery utilising either triplet (“UAG”) or quadruplet (“UAGA”) codons to direct incorporation at K201. Animals were grown in the presence of 1 mM PCK. Scale bars = 20 μm **(E)** Quantification of photocaged Cre activation for the lines depicted in (D). Animals were grown in the presence of 4 mM PCK for 48h. After uncaging, 30 animals were scored for expression of target gene. The mean of three independent experiments is shown. Error bars show the standard error of the mean. Significance is derived from P values of Welch’s t test.

We generated transgenic strains expressing the quadruplet-decoding system together with a construct expressing photocaged Cre, containing TAGA in the place of the codon for lysine K201 and a reporter construct to monitor Cre activity that we have previously described [25]. The reporter construct consists of a channelrhodopsin gene fused to red fluorescent mKate2, separated from its promoter by a transcriptional terminator sequence flanked by loxP sites (Figure 3A). Cre activate transcription of the reporter and results in red fluorescence. To visualise presence of the construct encoding photocaged Cre, we expressed it as part of an artificial operon together with the marker GFP (Figure 3A). All protein-coding genes were expressed from the *glr-1p* promoter that is active in *C. elegans* glutamatergic neurons [25, 33].

We grew age-synchronised populations of transgenic worms on NGM agar plates supplemented with 4 mM PCK for 48h, from the L1 larval stage. We then activated photocaged Cre by illumination with 365 nm light, and scored for expression of the target gene 24h after activation. For both quadruplet tRNA variants, we observed animals that showed clear expression of ChR2::mKate2 in neurons expressing photocaged Cre. In contrast, we saw no ChR2::mKate2 in animals grown on PCK but not subjected to UV illumination (Figure 3C). When we quantified Cre activation frequency as a measure of the production of photocaged Cre and thus PCK incorporation, we found that the most efficient tRNA variant tRNA(M15/UAGA-1)_UCUA_ performed equally well as the triplet variant tRNA(C15)_CUA_, with over 70% of animals showing expression of the target gene ChR2::mKate2. The less efficient tRNA(M15/M7)_UCUA_ still showed optically triggered Cre activity in 30% of animals. The optimised quadruplet incorporation machinery thus performs with a level of efficiency close to an optimised UAG triplet decoding system.

### Quadruplet dependent incorporation of photocaged cysteine

In *C. elegans* the only photocaged amino acids that have been successfully incorporated into proteins are photocaged lysine and photocaged tyrosine [25, 34]. We decided to apply quadruplet decoding to a further ncAA, namely a photocaged cysteine (PCC), thus expanding the repertoire of photocaged amino acids for *C. elegans*. Cysteine is an attractive target for photo-caging, since key cysteine residues are present in many biologically important protein classes, including ubiquitin ligases, phosphatases, and proteases such as deubiquitinases and caspases.

We used cysteine caged with a nitropiperonyl group (Figure 4A), which has an absorption maximum close to 365 nm and is therefore more amenable to *in vivo* use than the alternative *ortho*-nitrobenzyl caging group whose optical removal can be challenging [35]. Nitropiperonyl PCC is recognised by PCCRS, a variant of PylRS and has previously been established in mammalian cultured cells. We introduced the previously described PCCRS mutations [35] into a PylRS gene from *Methanosarcina mazei*, optimised for expression in *C. elegans*. To improve incorporation efficiency, we again added the Smad4 nuclear export sequence to the N-terminus to generate NES::PCCRS.

**Figure 4.**
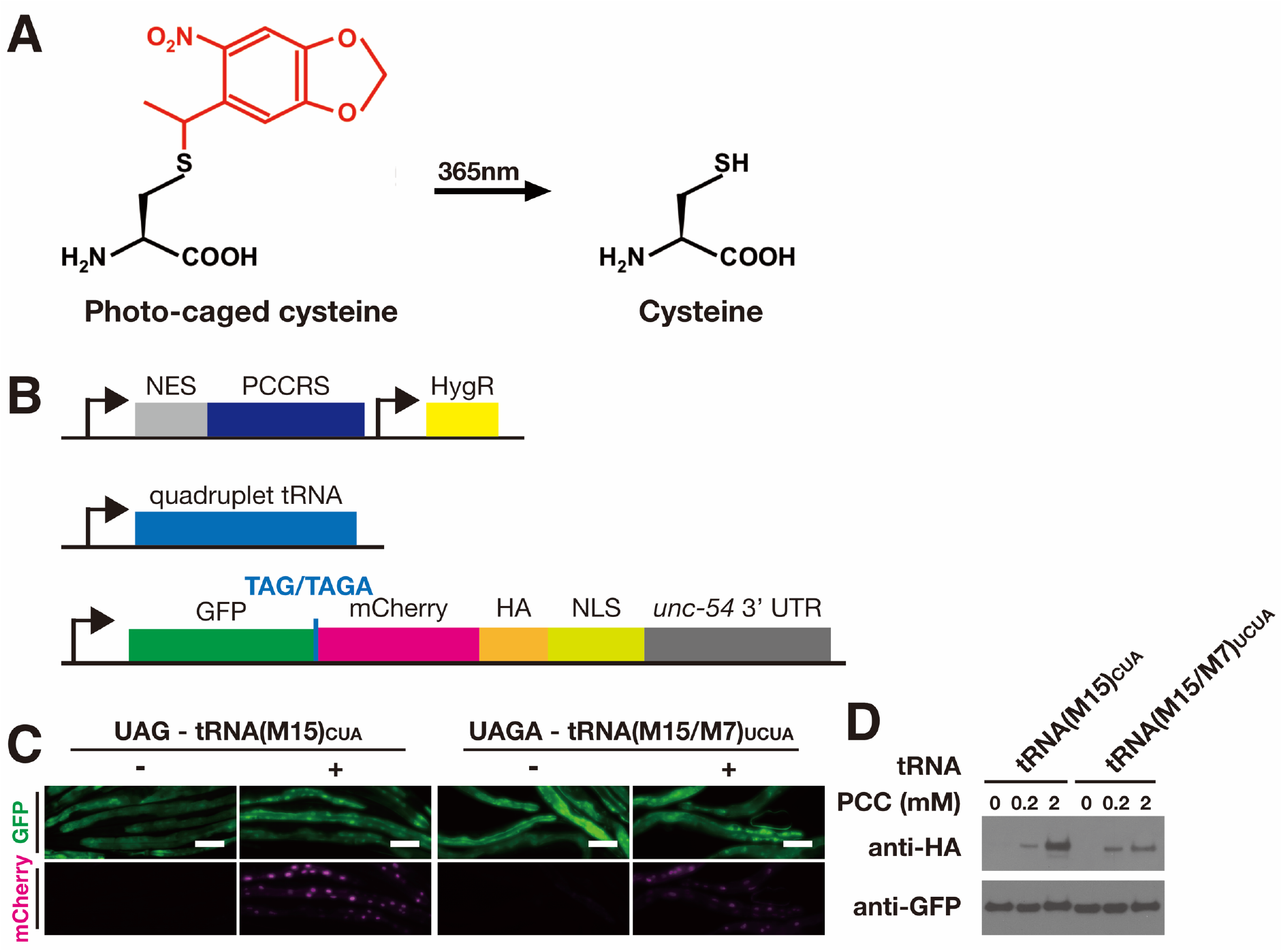
Incorporation of photocaged cysteine in *C. elegans*. **(A)** Genetic constructs for PCC-incorporating UAGA-decoding machinery, consisting of NES::PCCRS and quadruplet tRNA, and the fluorescent reporter GFP::TAGA::mCherry. **(B)** The nitropiperinyl caging group (red) of photocaged cysteine can be removed by 365-nm illumination. **(C)** Fluorescence microscopy images of transgenic lines expressing either the triplet tRNA(M15)_CUA_ or the quadruplet tRNA(M15/M7)_UCUA_, together with the aaRS and the fluorescent reporter as shown in (A). Animals were grown in the presence (“+”) or absence (“−”) of 2 mM PCC. Scale bar, 100 μm. **(D)** Western blots of the lines shown in (B) and grown in the absence or the presence of either 0.2 mM or 2 mM PCC. Full-length reporter protein was detected using anti-HA antibody, samples were normalised using anti-GFP antibody.

We created transgenic strains expressing NES::PCCRS together with either tRNA(M15)_CUA_ or tRNA(M15/M7)_UCUA_. To visualise incorporation, we co-expressed our fluorescent GFP::mCherry reporter with the two fluorescent proteins separated with either a UAG triplet codon to assay incorporation by the triplet tRNA(M15)_CUA_, or a UAGA quadruplet codon to assay incorporation by the quadruplet tRNA(M15/M7)_UCUA_ (Figure 4B).

When we grew transgenic animals on NGM agar plates supplemented with PCC, we observed strong red fluorescence appearing within 24h for both the triplet and the quadruplet system (Figure 4C). We further confirmed the identity of the full-length reporter protein by western blot against the C-terminal HA tag (Figure 4D). We thus show that PCC can be efficiently incorporated in *C. elegans* using either a UAG triplet or a UAGA quadruplet codon.

### Quadruplet codon directed incorporation of photocaged cysteine into Caspase-3 for optical control of apoptosis induction in *C. elegans*

We then decided to use the quadruplet system to incorporate PCC into the active site of human Caspase-3, a caspase that is routinely used for genetically controlled cell ablation in *C. elegans* [36]. By introducing PCC we aimed to optically control activity of the caspase and thus use light to ablate cells.

As a key executor of the apoptotic pathway, wild type Caspase-3 is synthesised as an inactive zymogen with its subunits arranged in such a way as to be sterically prevented from folding into the active conformation. The zymogen is converted into the active caspase by cleavage at internal proteolytic sites and subsequent rearrangement of its long and short subunits to form the active enzyme (Supplementary Figure 3A) [37, 38]. By switching the order of the subunits on the polypeptide chain, it is possible to express a constitutively active form of Caspase-3 that does not require proteolytic cleavage for activation (Supplementary Figure 3B) [37].

We based our design of photocaged caspase on the constitutively active form of Caspase-3, replacing the catalytically critical cysteine residue in the caspase active site with PCC. In this arrangement, activity of the enzyme would be blocked by the caging group on PCC and activity could later be restored by removing the caging group via 365 nm illumination (Figure 5A). The activation of photocaged caspase should then result in the death and removal of the targeted cell.

**Figure 5.**
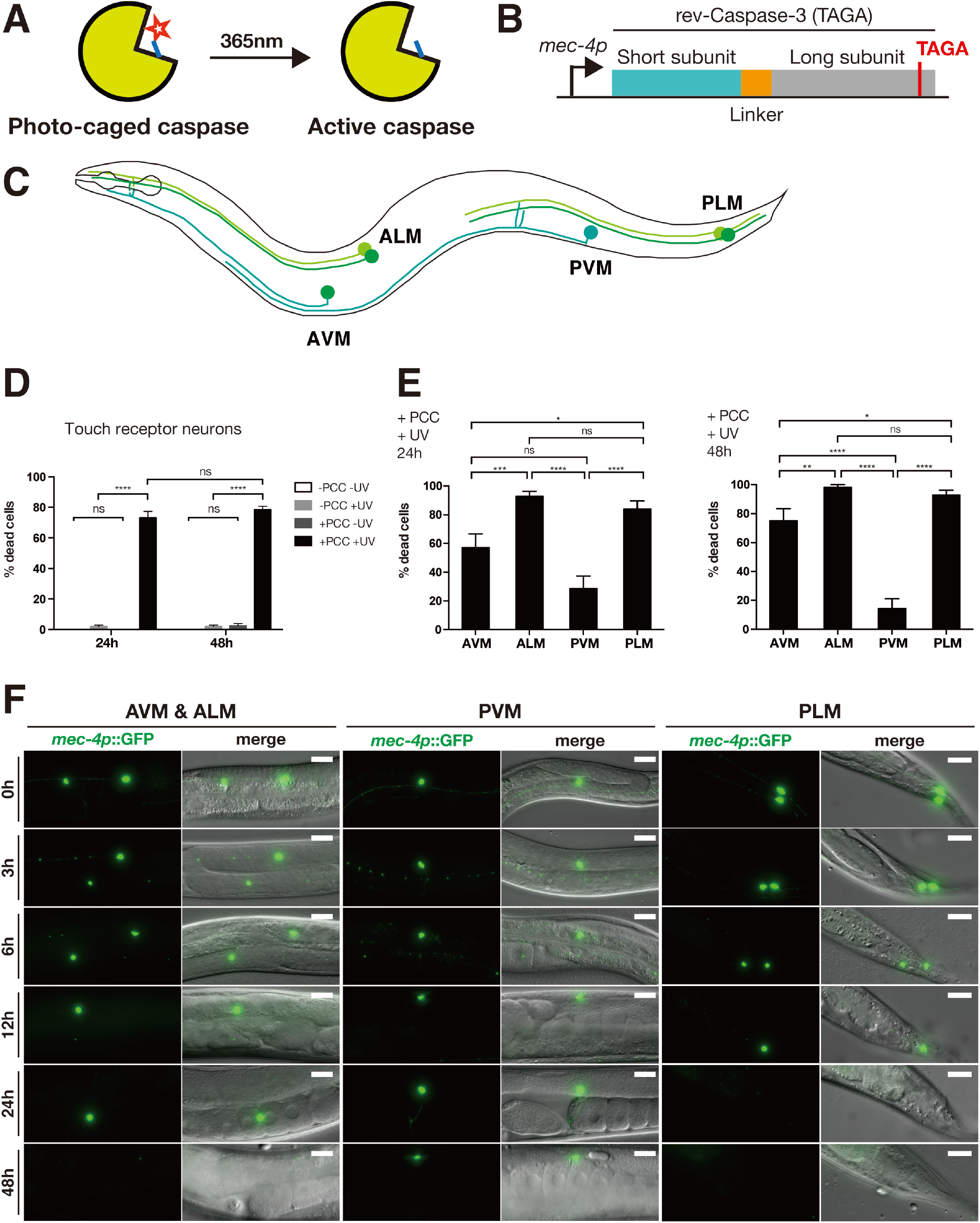
Optical control of caspase activity using quadruplet codon directed incorporation of photocaged cysteine. **(A)** Photocaged caspase with PCC (red star) in its active site is inactive, until it is uncaged by 365 nm light. **(B)** Genetic construct for expression of photocaged Caspase-3. Photocaged cysteine is inserted in place of the catalytic cysteine using a TAGA quadruplet codon. **(C)** Schematic of the six *C. elegans* touch receptor neurons (TRN). AVM and PVM are single cells, while ALM and PLM are bilaterally symmetric neurons pairs. **(D)** Percentages of TRN killed in grown in the presence (“+PCC”) or absence (“−PCC”) of photocaged cysteine and either uncaged by illumination with 365 nm (“+UV”) or kept in the dark (“−UV”). More than 25 animals were randomly picked for one treatment group and scored for missing neurons either 24h or 48h after the uncaging step. Shown are the average scores of all worms of each group. Error bars represent standard errors of mean. Significance between groups was determined by Friedman’s 2-way ANOVA and *post hoc* sign tests. **(E)** Percentage of disappeared cells by TRN type of the “+PCC +UV” animals assayed in (D) 24h (left graph) and 48h (right graph) post UV uncaging. Shown are the average scores of all worms of each group. Error bars represent standard errors of mean. Significance was determined by Fisher’s exact tests. **(F)** Fluorescent images of dying neurons 0, 3, 6, 12, 24, 48h after UV uncaging. GFP signals were superimposed onto DIC images to generate merged images. Scale bar = 20 μm.

We generated transgenic strains expressing the quadruplet incorporation machinery for PCC, consisting of NES::PCCRS and tRNA(M15/M7)_UCUA_ together with a construct encoding for photocaged caspase, which contained a TAGA quadruplet codon in place of the catalytic cysteine (Figure 5B). We targeted expression of NES::PCCRS and photocaged caspase to the *C. elegans* touch receptor neurons (TRN) using the *mec-4p* promoter (Figure 5B,C) [39]. We chose the TRNs, a neuron class consisting of 6 cells, since they are easy to distinguish and score under a microscope. We generated transgenic lines in the CZ10175 strain, which carries a genomically integrated transcriptional *mec-4p*::*gfp* fusion, labelling the TRN neurons, but is otherwise wild type. The GFP fluorescence allowed us to easily score for presence or absence of neurons to quantify removal of cells upon photocaged caspase activation.

We grew age-synchronised populations of transgenic worms on NGM agar plates supplemented with 2 mM PCC from the L1 larval stage for 3 days until the late L4 larval stage. To activate photocaged caspase, we then uncaged PCC using 365 nm illumination and scored the number of neurons 24h after uncaging. We found that animals grown on PCC and treated with UV had lost an average of 73% of TRNs (Figure 5D). This number did not increase significantly after 48h, indicating that the majority of cell removal due to the optically induced caspase activity occurred within 24h after activation.

Interestingly, we found that among the TRNs some cell types were removed more effectively than others, with the ALM and the PLM neuron pairs showing the greatest proclivity for photocaged caspase induced cell ablation. 48h post activation, almost 100% of ALM and PLM cells had disappeared. This was in contrast to the AVM neuron (75% disappeared after 48h) and especially the PVM neuron (14% disappeared) (Figure 5E).

When we examined cells at different time points after caspase activation, we observed changes in cell morphology consisting of fragmentation of cell processes as a first sign of initiated cell degeneration, followed by the cells taking on a spherical appearance and finally complete disappearance (Figure 5F).

We could therefore demonstrate that the improved quadruplet incorporation machinery allows for the use of PCC to produce sufficient photocaged caspase for optically induced caspase dependent cell ablation in *C. elegans* neurons.

### Discussion

The site-specific incorporation of ncAA in multicellular organisms was until now limited to the use of reassigned UAG stop codons. We report here the first instance of genetic code expansion using quadruplet decoding in an animal. Our system reaches incorporation efficiencies for a quadruplet codon approaching those observed with optimised UAG triplet decoding tRNAs and surpassing those of a non-optimised UAG triplet system. We achieve this by establishing the use of hybrid tRNA variants, which combine scaffolds optimised for use in eukaryotic cells and anticodon loops evolved for decoding of quadruplet codons.

We observed striking efficiency improvements for all four tested hybrid tRNAs compared to their parent molecule scaffolds and anticodon loops. Importantly, this suggests that it may be possible to treat anticodon loops and scaffolds as independent biological parts that can be combined to allow the creation of novel tRNA hybrids. An immediate benefit would be the possibility of independently evolving such parts before easily recombining the newly evolved structural features into tRNA hybrids. This is an attractive prospect, given that several mutually orthogonal and efficient aminoacyl-tRNA-synthetase / tRNA pairs based on the PylRS/tRNA(Pyl) system have recently been developed for bacteria and mammalian cells [40, 41], albeit they have not yet been established in animals. In order to utilise such mutually orthogonal pairs, it will be necessary to have access to sufficient coding space to allow for the possibility of incorporating more than one and up to several ncAAs independently within the same cell. Since the identity elements of these newly developed orthogonal PylRS/tRNA(Pyl) pairs lie within the scaffold region and outside the anticodon loop it should be possible to use them for the construction of new orthogonal quadruplet hybrid tRNAs and thus facilitate the expansion of the coding space in cells and organisms. Likewise, it should be possible to evolve or rationally redesign scaffolds using a UAG triplet anticodon and then combine such improvements with independently evolved specialised quadruplet anticodon loops.

Importantly, we were also able to show that improvements in anticodon loops resulting in increased quadruplet decoding efficiency are to some extent independent of the anticodon itself. We found that the M7 anticodon loop with a UCUA anticodon could efficiently decode the UAGA quadruplet, even though the M7 anticodon loop was originally evolved for the UCCU anticodon and decoding of the quadruplet AGGA [16]. This suggests the possibility of directly utilising the hybrid tRNAs presented here for decoding other quadruplet codons without the need for further optimisation, by simply replacing the anticodon.

We show that it is possible to achieve quadruplet incorporation efficiencies that, while still below those observed with an optimised UAG triplet system, can easily produce sufficient amounts of protein for *in vivo* applications. Indeed, we find that in a direct comparison using photocaged Cre recombinase, our quadruplet system performs as efficiently as an optimised UAG triplet system.

We were able to use quadruplet decoding in conjunction with photocaged cysteine to express photocaged caspase as a novel tool for cell ablation in *C. elegans*. Cell ablation is a widely used approach to uncover the function of cells within the organismal context. A number of cell-ablation methods have been developed in *C. elegans* and other systems, namely laser ablation [42], expression of toxic dominant mutations in ion channels such as *mec-4(d)* [43] or *trp-4(d)* [44] and optogenetic approaches using singlet oxygen generators such as MiniSOG [45]. Laser ablation, while allowing the targeting of individual cells, can be technically challenging, and is in most cases not to perform without affecting surrounding tissues in adult worms with a fully developed nervous system. The expression of toxic proteins is dependent on the availability of cell specific promoters and does not offer temporal precision. Singlet oxygen generators may require prolonged illumination and are not compatible with imaging approaches utilising the same visible wavelengths used to activate singlet oxygen production. The use of an optically controlled caspase allows a temporally controlled removal of cells, while the use of a UV responsive photocage allows for the possibility of using visible wavelengths for imaging or a control of additional optogenetic tools. Furthermore, short, low-powered UV pulses delivered using a microscope-mounted laser may allow the targeted removal of individual cells without affecting surrounding tissues. We have previously demonstrated the feasibility of targeted laser uncaging in *C. elegans* for the activation of photocaged Cre recombinase [25].

The repurposing of triplet stop codons as sense codons for genetic code expansion is accompanied by the possible misincorporation of ncAA at endogenous stop codons [46], which may lead to extension of native protein products and increased cellular stress. Such misincorporation events are especially undesirable for experimental approaches such as protein labelling or crosslinking, where it may contribute to non-specific background signals. The UCUA quadruplet anticodon used in our study, which decodes UAGA, may help to alleviate such undesired misincorporation effects as it was previously shown to result in significantly lower levels of cross-decoding at UAGN codons other than UAGA [18]. The routine use of quadruplet codons for genetic code expansion may therefore be highly desirable, even when incorporating only a single ncAA. Our improved quadruplet decoding system makes the routine use of quadruplets possible. In fact, since it outperforms the standard wild type PylRS / tRNA(Pyl)_CUA_ triplet pair that is currently used for most genetic code expansion work in animals and cultured cells, a switch to quadruplets will not come at the price of reduced efficiency for most labs.

In summary, we have succeeded in adding quadruplet codons to the genetic coding repertoire of a multicellular organism, by introducing the use of hybrid tRNAs combining optimised anticodon loops and scaffolds. The system we have developed reaches efficiency levels for the site-specific incorporation of a ncAA that come close to those observed for UAG triplet codons. Using photocaged amino acids encoded by quadruplet codons, we were able to express photocaged Cre recombinase and photocaged caspase allowing us to optically control gene expression and to induce cell death in *C. elegans* neurons. We therefore demonstrate that quadruplet codons are a viable alternative to triplet codons for genetic code expansion. We expect that the improvements we have demonstrated will be applicable in single- and multicellular eukaryotic systems beyond *C. elegans*.

## Materials and Methods

### Plasmids

All expression plasmids were constructed from pENTR plasmids using Gateway cloning (Thermo Fisher Scientific). DNA fragments were either PCR-amplified with oligonucleotide primers (Sigma-Aldrich and IDT) or ordered as gBlocks from IDT and cloned into Gateway pENTR plasmids. Synthetic genes were optimised for *C. elegans* expression [47]. Expression constructs were assembled from pENTR plasmids using Gateway cloning (Thermo Fisher Scientific). All plasmids are described in Supplementary Table 3.

### *C. elegans* strains and maintenance

Strains were maintained under standard conditions unless otherwise indicated [48, 49]. Transgenic lines were generated by biolistic bombardment into the N2, *smg-6(ok1794)* or CZ10175 genetic backgrounds using hygromycin B (Formedium) as a selection marker [31, 50]. Details are listed in Supplementary Table 2. Transgenic lines were maintained on hygromycin B. No hygromycin B was added to plates containing ncAA.

### Feeding of non-canonical amino acids

Photocaged lysine (PCK) and photocaged cysteine (PCC) were custom synthesized by ChiroBlock GmbH. To prepare PCK/PCC supplemented NGM plates, the ncAA powder was first dissolved in a small volume of 0.02M HCl and then mixed into molten NGM agar with equimolar amounts of NaOH added to the NGM agar for neutralisation [50]. Worms were age synchronised by bleaching [49] and added to ncAA plates as L1 larvae. Freeze-dried OP50 (LabTIE) was reconstituted according to the manufacturer’s instructions and added as food. The ncAA and ncAA plates were stored in the dark, and animals on ncAA were grown in the dark.

### Worm lysis and western blotting

Lysis was performed as previously described [25]. Samples were run on precast Bolt 4-12% Bis-Tris Plus polyacrylamide gels (Thermo Fisher Scientific) in Bolt MES SDS running buffer at 200V for 22 minutes. Proteins were transferred onto a nitrocellulose membrane using an iBlot2 device (Thermo Fisher Scientific). After transfer, the membrane was blocked for 1 h at room temperature using PBST (PBS + 0.1% Tween-20) supplemented with 5% milk powder. Incubation with primary antibodies was carried out at 4°C overnight. Blots were then washed 6 times for 5 minutes in PBST + 5% milk powder. Blots were then incubated with secondary antibodies diluted in PBST + 5% milk powder for 1h at room temperature, followed by 3 washes of 5 minutes using PBST + 5% milk powder and one wash with PBS. Primary antibodies used were rat anti-HA (clone 3F10, Roche) at a dilution of 1:3000 for blots exposed to film, 1:1500 for quantitative western blots, and mouse anti-GFP (clones 7.1 and 13.1, Roche) at a dilution of 1:3000 for blots exposed to film, 1:5,000 for quantitative western blots. Secondary antibodies were goat anti-rat IgG(H+L)-HRP (Thermo Fisher Scientific) at a dilution of 1:5000, and horse anti-mouse IgG-HRP (Cell Signaling Technology) at a dilution of 1:5000. All dilutions were made in PBST + 5% milk powder.

Pierce ECL Western Blotting Substrate (Thermo Fisher Scientific) or SuperSignal West Femto Maximum Sensitivity Substrate (Thermo Fisher Scientific) were used as detection agents. For quantitative western blots, chemiluminescence was measured using a C-DiGit Blot Scanner (LI-COR) and band intensities analysed using ImageStudio software.

### Imaging

All imaging was performed using a Zeiss Axio Imager M2 microscope and images processed using ZEN software (Zeiss) and ImageJ software. For imaging, animals were anaesthetised in a drop of M9 with 5mM levamisole (Sigma-Aldrich) or 25mM NaN_3_ (Merck) and mounted on a 3% agarose pad cast on a glass slide.

### Uncaging of PCK for Cre activation

Synchronized L1 larvae were grown for 48h on NGM plates supplemented with 4 mM PCK or without PCK, then washed onto unseeded NGM plates and illuminated in a 365 nm CL-1000L crosslinker (UVP) at 5 mW/cm^2^ for 5 minutes as previously described [50]. After uncaging FUDR was added to a final concentration of 400 μM to prevent hatching of F1 progeny and thus aid scoring of animals expressing the target gene.

48 hours after uncaging, animals were scored for expression of the target gene using a Leica M165FC fluorescence dissection scope under a 2.0x objective. Each plate was independently and blindly scored by two people and the mean of both counts was used. All experiments were performed in triplicate. Significance tests were carried out using Welch’s t test as a pairwise comparison between each condition at each concentration using Prism6 software.

### Uncaging of PCC for Caspase-3 activation and apoptosis assay

To facilitate scoring and imaging, caspase expressing strains were constructed in the CZ10175 strain background, which contains a genomically integrated *mec-4p::gfp* transcriptional fusion to express GFP in the touch receptor neurons. Synchronized L1 larvae were grown for 48h on NGM plates with either 0 or 2 mM PCC, before being washed onto unseeded 6cm NGM plates and illuminated in a CL-1000L UV crosslinker (UVP) at 365 nm, 5 mW/cm^2^ for 5 minutes.

To document morphological changes upon caspase activation, worms were mounted and imaged at 0, 3, 6, 12, 24, 48h after uncaging. To quantify ablation efficiency, worms were randomly picked at 24 and 48h after uncaging and scored for the presence or absence of target neurons, using a Zeiss Axio Imager M2. Each condition was scored blind and repeated in duplicate. Significance tests of the percentage of disappeared cells among all TRNs were carried out using Friedman’s 2-way ANOVA by ranks and then sign tests as a pairwise comparison between each condition at each concentration, by SPSS Statistics 23. Significance tests of the cell disappearance by TRN type were carried out between each condition using Fisher’s exact tests, by Prism6 software.

## Acknowledgements

We thank Maria Doitsidou, Jack O’Shea, Emanuel Busch, and members of the Greiss, Doitsidou and Busch labs for helpful suggestions on the manuscript. We thank the European Research Council (ERC-StG-679990), the Muir Maxwell Epilepsy Centre, the Royal Society, and the Wellcome-Trust University of Edinburgh Institutional Strategic Support Fund ISS2 for funding to S.G., the University of Edinburgh for an Edinburgh Global Award and Principal’s Career Development PhD studentship to Z.X. Strain CZ10175 was provided by the Caenorhabditis Genetics Centre for strains, funded by NIH Office of Research Infrastructure Programs (P40OD010440)

## References

1. Crick, F.H.C., et al., General nature of the genetic code for proteins. Nature, 1961. 192: p. 1227–32.

2. Chambers, I., et al., The structure of the mouse glutathione peroxidase gene: the selenocysteine in the active site is encoded by the ‘termination’ codon, TGA. EMBO J, 1986. 5(6): p. 1221–7.

3. Zinoni, F., et al., Nucleotide sequence and expression of the selenocysteine-containing polypeptide of formate dehydrogenase (formate-hydrogen-lyase-linked) from Escherichia coli. Proc Natl Acad Sci U S A, 1986. 83(13): p. 4650–4.

4. Hao, B., et al., A new UAG-encoded residue in the structure of a methanogen methyltransferase. Science, 2002. 296(5572): p. 1462–6.

5. Srinivasan, G., C.M. James, and J.A. Krzycki, Pyrrolysine encoded by UAG in Archaea: charging of a UAG-decoding specialized tRNA. Science, 2002. 296(5572): p. 1459–62.

6. Krauskopf, K. and K. Lang, Increasing the chemical space of proteins in living cells via genetic code expansion. Current Opinion in Chemical Biology, 2020. 58: p. 112–120.

7. Wang, L., et al., Expanding the Genetic Code of Escherichia coli. Science, 2001. 292: p. 498–500.

8. Chin, J.W., et al., An Expanded Eukaryotic Genetic Code. Science, 2003. 301(5635): p. 964–7.

9. Hino, N., et al., Protein photo-cross-linking in mammalian cells by site-specific incorporation of a photoreactive amino acid. Nat Methods., 2005. 2(3): p. 201–6.

10. Greiss, S. and J.W. Chin, Expanding the genetic code of an animal. J Am Chem Soc, 2011. 133(36): p. 14196–9.

11. Bianco, A., et al., Expanding the genetic code of Drosophila melanogaster. Nat Chem Biol., 2012. 8(9): p. 748–50.

12. Chen, Y., et al., Heritable expansion of the genetic code in mouse and zebrafish. Cell Res., 2017. 27(2): p. 294–297.

13. Ostrov, N., et al., Design, synthesis, and testing toward a 57-codon genome. Science, 2016. 353(6301): p. 819–22.

14. Wang, K., et al., Defining synonymous codon compression schemes by genome recoding. Nature, 2016. 539(7627): p. 59–64.

15. Anderson, J.C., et al., An expanded genetic code with a functional quadruplet codon. Proc Natl Acad Sci U S A, 2004. 101(20): p. 7566–71.

16. Niu, W., P.G. Schultz, and J. Guo, An expanded genetic code in mammalian cells with a functional quadruplet codon. ACS Chem Biol, 2013. 8(7): p. 1640–5.

17. Chatterjee, A., et al., A Bacterial Strain with a Unique Quadruplet Codon Specifying Non-native Amino Acids. ChemBioChem, 2014. 15(12): p. 1782–1786.

18. Wang, N., et al., Systematic Evolution and Study of UAGN Decoding tRNAs in a Genomically Recoded Bacteria. Scientific Reports, 2016. 6(1).

19. Taki, M., J. Matsushita, and M. Sisido, Expanding the Genetic Code in a Mammalian Cell Line by the Introduction of Four-Base Codon/Anticodon Pairs. ChemBioChem, 2006. 7(3): p. 425–428.

20. de la Torre, D. and J.W. Chin, Reprogramming the genetic code. Nature Reviews Genetics, 2020. 22(3): p. 169–184.

21. Wang, K., et al., Optimized orthogonal translation of unnatural amino acids enables spontaneous protein double-labelling and FRET. Nat Chem, 2014. 6(5): p. 393–403.

22. Neumann, H., et al., Encoding multiple unnatural amino acids via evolution of a quadruplet-decoding ribosome. Nature, 2010. 464(7287): p. 441–444.

23. Chen, Y., et al., Controlling the Replication of a Genomically Recoded HIV-1 with a Functional Quadruplet Codon in Mammalian Cells. ACS Synthetic Biology, 2018. 7(6): p. 1612–1617.

24. Serfling, R., et al., Designer tRNAs for efficient incorporation of non-canonical amino acids by the pyrrolysine system in mammalian cells. Nucleic Acids Res, 2017. 1.

25. Davis, L., et al., Precise optical control of gene expression in C. elegans using genetic code expansion and Cre recombinase, U.o. Edinburgh, Editor. 2020: http://www.bioRxiv.org.

26. Gautier, A., et al., Genetically Encoded Photocontrol of Protein Localization in Mammalian Cells. J Am Chem Soc, 2010: p. 4086–8.

27. Watanabe, M., et al., Regulation of intracellular dynamics of Smad4 by its leucine-rich nuclear export signal. EMBO Rep., 2000. 1(2): p. 176–82.

28. Nikić, I., et al., Debugging Eukaryotic Genetic Code Expansion for Site-Specific Click-PAINT Super-Resolution Microscopy. Angewandte Chemie International Edition, 2016. 55(52): p. 16172–16176.

29. Parrish, A.R., et al., Expanding the genetic code of Caenorhabditis elegans using bacterial aminoacyl-tRNA synthetase/tRNA pairs. ACS Chem Biol, 2012. 7(7): p. 1292–302.

30. Yochem, J., T. Gu, and M. Han, A New Marker for Mosaic Analysis in Caenorhabditis elegans Indicates a Fusion Between hyp6 and hyp7, Two Major Components of the Hypodermis. Genetics, 1998. 149(3): p. 1323–34.

31. Radman, I., S. Greiss, and J.W. Chin, Efficient and rapid C. elegans transgenesis by bombardment and hygromycin B selection. PLoS One, 2013. 8(10): p. e76019.

32. Luo, J., et al., Genetically encoded optical activation of DNA recombination in human cells. Chem Commun (Camb), 2016. 52(55): p. 8529–32.

33. Maricq, A.V., et al., Mechanosensory signalling in C. elegans mediated by the GLR-1 glutamate receptor. Nature, 1995. 378(6552): p. 78–81.

34. O’Shea, J.M., et al., Generation of photocaged nanobodies for in vivo applications using genetic code expansion and computationally guided protein engineering, U.o. Edinburgh, Editor. 2021: http://www.bioRxiv.org.

35. Nguyen, D.P., et al., Genetic encoding of photocaged cysteine allows photoactivation of TEV protease in live mammalian cells. J Am Chem Soc, 2014. 136(6): p. 2240–3.

36. Chelur, D.S. and M. Chalfie, Targeted cell killing by reconstituted caspases. Proc Natl Acad Sci U S A, 2007. 104(7): p. 2283–8.

37. Srinivasula, S.M., et al., Generation of constitutively active recombinant caspases-3 and −6 by rearrangement of their subunits. J Biol Chem 1998. 273(17): p. 10107–11.

38. Slee, E.A., et al., Ordering the cytochrome c-initiated caspase cascade: hierarchical activation of caspases−2, −3, −6, −7, −8, and −10 in a caspase-9-dependent manner. J Cell Biol., 1999. 144(2): p. 281–92.

39. Lai, C.C., et al., Sequence and Transmembrane Topology of MEC-4, an Ion Channel Subunit Required for Mechanotransduction in Caenorhabditis elegans. J Cell Biol, 1996. 133(5): p. 1071–81.

40. Meineke, B., et al., Methanomethylophilus alvus Mx1201 Provides Basis for Mutual Orthogonal Pyrrolysyl tRNA/Aminoacyl-tRNA Synthetase Pairs in Mammalian Cells. ACS Chemical Biology, 2018. 13(11): p. 3087–3096.

41. Willis, J.C.W. and J.W. Chin, Mutually orthogonal pyrrolysyl-tRNA synthetase/tRNA pairs. Nat Chem, 2018.

42. Fang-Yen, C., et al., Laser Microsurgery in Caenorhabditis elegans. Methods Cell Biol., 2012. 107: p. 177–206.

43. Harbinder, S., et al., Genetically Targeted Cell Disruption in Caenorhabditis Elegans. Proc Natl Acad Sci U S A, 1997. 94(24): p. 13128–33.

44. Nagarajan, A., et al., Progressive Degeneration of Dopaminergic Neurons through TRP Channel-Induced Cell Death. Journal of Neuroscience, 2014. 34(17): p. 5738–5746.

45. Xu, S. and A.D. Chisholm, Highly efficient optogenetic cell ablation in C. elegans using membrane-targeted miniSOG. Scientific Reports, 2016. 6(1).

46. Aerni, H.R., et al., Revealing the amino acid composition of proteins within an expanded genetic code. Nucleic Acids Research, 2015. 43(2): p. e8–e8.

47. Redemann, S., et al., Codon adaptation-based control of protein expression in C. elegans. Nat Methods, 2011. 8(3): p. 250–4.

48. Brenner, S., The genetics of Caenorhabditis elegans. Genetics, 1974. 77(1): p. 71–94.

49. Stiernagle, T., Maintenance of C. elegans. WormBook, 2006: p. 1–11.

50. Davis, L. and S. Greiss, Genetic Encoding of Unnatural Amino Acids in C. elegans, in Noncanonical Amino Acids E.A. Lemke, Editor. 2018, Humana Press: New York. p. 389–408.

