## Supplementary Figures and Tables for "Using a Quadruplet Codon to Expand the Genetic Code of an Animal"

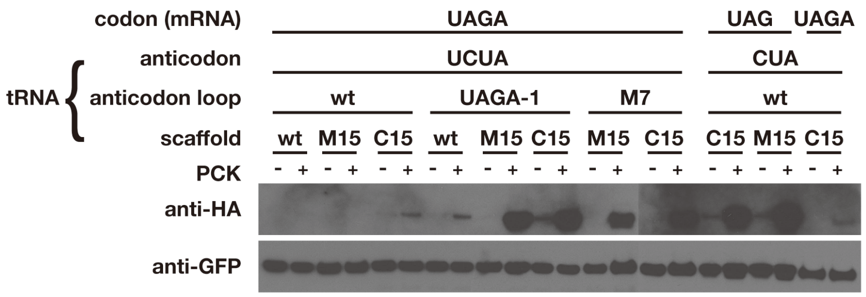


**Supplementary Figure 1: Extended exposure of western blot shown in Figure 2D.**

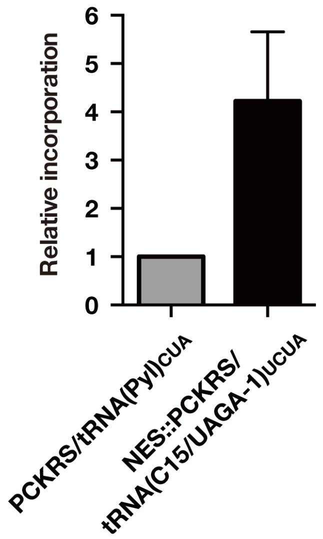


**Supplementary Figure 2: Quantitative western blots comparing quadruplet to non-optimised triplet decoding**.

Quantitative western blots of the most efficient quadruplet system (consisting of NES::PCKRS and tRNA(C15/UAGA-1)$_{UCUA}$) compared to the non-optimised triplet system (consisting of PCKRS and tRNA(Pyl)$_{CUA}$). Relative incorporation was determined by dividing the intensity of the full-length product bands for each experiment with the mean intensity of the truncated GFP products. Incorporation was performed in the presence of 1mM PCK. The full-length product was detected using anti-HA antibody, the truncated product was detected using anti-GFP antibody. Two independent lines were assayed for each condition. For PCKRS/tRNA(Pyl)$_{CUA}$ each line was independently grown on PCK twice, NES::PCKRS/tRNA(C15/UAGA-1)$_{UCUA}$ was grown independently on PCK three times. Each sample was blotted and measured twice. Error bar shows standard error of the mean.

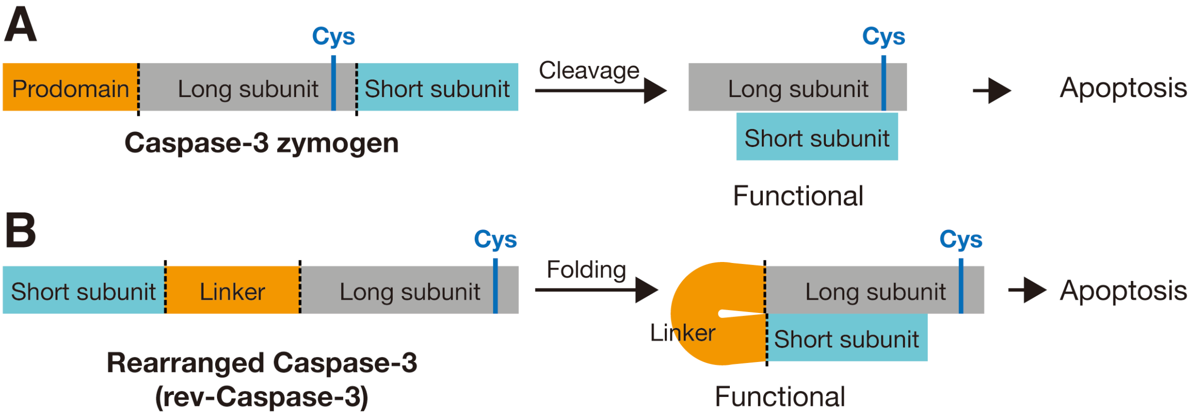


**Supplementary Figure 3: Schematic of Caspase-3 activation.**
**(A)** From N- to C-terminus, human Caspase-3 zymogen consists of a prodomain (orange), long subunit (grey) and short subunit (cyan), all separated by caspase cleavage sites (dash line). Cleavage removes the prodomain and enables assembly of the long and short subunits to form active caspase. **(B)** Caspases can be expressed in a constitutively active form by reversing the subunit order in the polypeptide [1].

### Supplementary Table 1 - Sequence of tRNA variants

The codons for incorporation are marked in red and bold. Nucleotides of the tRNA variants different from tRNA(Pyl)$_{UCUA}$ are underlined.

| Name | Sequence |
|---|---|
| tRNA(Pyl)$_{UCUA}$ | GGAAACCTGATCATGTAGATCGAATGGACT**TCTA**AATCCGTTCAGCCGGGTTAGATTCCCGGGGTTTCCG |
| tRNA(M15)$_{CUA}$ | GGAAACCTGGTCAGGGAGACCGAACGGACT**CTA**AATCCGTTCAGCCGGGTTcGATTCCCGGGGTTTCCG |
| tRNA(C15)$_{CUA}$ | GGGAGAGTGGCCAAGGTGGCCGTGTTGACT**CTA**AATCAACACAGGGGGGTTCGATTCCCCCCTCTCCCG |
| tRNA(M15/M7)$_{UCUA}$ | GGAAACCTGGTCAGGGAGACCGAAGGGGCT**TCTA**ATCCGCTTCAGCCGGGTTCGATTCCCGGGGTTTCCG |
| tRNA(M15)$_{UCUA}$ | GGAAACCTGGTCAGGGAGACCGAACGGACT**TCTA**AATCCGTTCAGCCGGGTTCGATTCCCGGGGTTTCCG |
| tRNA(C15/M7)$_{UCUA}$ | GGGAGAGTGGCCAAGGTGGCCGTGGGGGCT**TCTA**ATCCGCCACAGGGGGGTTCGATTCCCCCCTCTCCCG |
| tRNA(C15)$_{UCUA}$ | GGGAGAGTGGCCAAGGTGGCCGTGTTGACT**TCTA**AATCAACACAGGGGGGTTCGATTCCCCCCTCTCCCG |
| tRNA(Pyl/UAGA-1)$_{UCUA}$ | GGAAACCTGATCATGTAGATCGAATGGGCT**TCTA**ATCCTGTTCAGCCGGGTTAGATTCCCGGGGTTTCCG |
| tRNA(M15/UAGA-1)$_{UCUA}$ | GGAAACCTGGTCAGGGAGACCGAACGGGCT**TCTA**ATCCTGTTCAGCCGGGTTCGATTCCCGGGGTTTCCG |
| tRNA(C15/UAGA-1)$_{UCUA}$ | GGGAGAGTGGCCAAGGTGGCCGTGTGGGCT**TCTA**ATCCTACACAGGGGGGTTCGATTCCCCCCTCTCCCG |

### Supplementary Table 2 - Transgenic *C. elegans* Strains

| Name | Description | Plasmid 1 | Plasmid 2 | Plasmid 3 | Plasmid 4 |
|---|---|---|---|---|---|
| SGR46 | *greEx33[sur-5p::Hu smad-4 NES::PCKRS; rpr-1p::tRNA(C15)*$_{CUA}$*; rps-0p::GFP(TAG)::mCherry::HA::egl-13 NLS]* | SE170 | SE150 | SG88 | / |
| SGR48 | *greEx35[sur-5p::Hu smad-4 NES::PCKRS; rpr-1p::tRNA(C15)*$_{CUA}$*; rps-0p::GFP(TAG)::mCherry::HA::egl-13 NLS]* | SE170 | SE150 | SG88 | / |
| SGR51 | *greEx38[glr-1p::Hu smad-4 NES::PCKRS; rpr-1p::tRNA(C15)*$_{CUA}$*; glr-1p::Cre K201TAG::SL2::GFP; glr-1p::loxP::B-gal terminator + loxP::ChR2::mKate2]* | SE174 | SE150 | ZS11 | IR361 |
| SGR69 | *greEx55[sur-5p::Hu smad-4 NES::PCKRS; rpr-1p::tRNA(M15)*$_{CUA}$*; rps-0p::GFP(TAG)::mCherry::HA::egl-13 NLS]* | SE170 | SE149 | SG88 | / |
| SGR70 | *greEx56[sur-5p::Hu smad-4 NES::PCKRS; rpr-1p::tRNA(M15)*$_{CUA}$*; rps-0p::GFP(TAG)::mCherry::HA::egl-13 NLS]* | SE170 | SE149 | SG88 | / |
| SGR71 | *greEx57[sur-5p::Hu smad-4 NES::PCKRS; rpr-1p::tRNA(M15)*$_{UCUA}$*; rps-0p::GFP(TAGA)::mCherry::HA::egl-13 NLS]* | SE170 | SE292 | KB184 | / |
| SGR72 | *greEx58[sur-5p::Hu smad-4 NES::PCKRS; rpr-1p::tRNA(M15)*$_{UCUA}$*; rps-0p::GFP(TAGA)::mCherry::HA::egl-13 NLS]* | SE170 | SE292 | KB184 | / |

| | | | | | |
|---|---|---|---|---|---|
| SGR73 | *greEx59[sur-5p::Hu smad-4 NES::PCKRS; rpr-1p::tRNA(C15)$_{UCUA}$; rps-0p::GFP(TAGA)::mCherry::HA::egl-13 NLS]* | SE170 | SE296 | KB184 | / |
| SGR74 | *greEx60[sur-5p::Hu smad-4 NES::PCKRS; rpr-1p::tRNA(C15)$_{UCUA}$; rps-0p::GFP(TAGA)::mCherry::HA::egl-13 NLS]* | SE170 | SE296 | KB184 | / |
| SGR75 | *greEx61[sur-5p::Hu smad-4 NES::PCKRS; rpr-1p::tRNA(Pyl/UAGA-1)$_{UCUA}$; rps-0p::GFP(TAGA)::mCherry::HA::egl-13 NLS]* | SE170 | SE301 | KB184 | / |
| SGR76 | *greEx62[sur-5p::Hu smad-4 NES::PCKRS; rpr-1p::tRNA(Pyl/UAGA-1)$_{UCUA}$; rps-0p::GFP(TAGA)::mCherry::HA::egl-13 NLS]* | SE170 | SE301 | KB184 | / |
| SGR77 | *greEx63[sur-5p::Hu smad-4 NES::PCKRS; rpr-1p::tRNA(M15/UAGA-1)$_{UCUA}$; rps-0p::GFP(TAGA)::mCherry::HA::egl-13 NLS]* | SE170 | SE302 | KB184 | / |
| SGR78 | *greEx64[sur-5p::Hu smad-4 NES::PCKRS; rpr-1p::tRNA(M15/UAGA-1)$_{UCUA}$; rps-0p::GFP(TAGA)::mCherry::HA::egl-13 NLS]* | SE170 | SE302 | KB184 | / |
| SGR79 | *greEx65[sur-5p::Hu smad-4 NES::PCKRS; rpr-1p::tRNA(C15/UAGA-1)$_{UCUA}$; rps-0p::GFP(TAGA)::mCherry::HA::egl-13 NLS]* | SE170 | SE303 | KB184 | / |
| SGR80 | *greEx66[sur-5p::Hu smad-4 NES::PCKRS; rpr-1p::tRNA(C15/UAGA-1)$_{UCUA}$; rps-0p::GFP(TAGA)::mCherry::HA::egl-13 NLS]* | SE170 | SE303 | KB184 | / |
| SGR81 | *greEx67[sur-5p::Hu smad-4 NES::PCKRS; rpr-1p::tRNA(Pyl)$_{UCUA}$; rps-0p::GFP(TAGA)::mCherry::HA::egl-13 NLS]* | SE170 | SE304 | KB184 | / |
| SGR82 | *greEx68[sur-5p::Hu smad-4 NES::PCKRS; rpr-1p::tRNA(Pyl)$_{UCUA}$; rps-0p::GFP(TAGA)::mCherry::HA::egl-13 NLS]* | SE170 | SE304 | KB184 | / |
| SGR83 | *greEx69[sur-5p::Hu smad-4 NES::PCKRS; rpr-1p::tRNA(C15)$_{CUA}$; rps-0p::GFP(TAGA)::mCherry::HA::egl-13 NLS]* | SE170 | SE150 | KB184 | / |
| SGR84 | *greEx70[sur-5p::Hu smad-4 NES::PCKRS; rpr-1p::tRNA(C15)$_{CUA}$; rps-0p::GFP(TAGA)::mCherry::HA::egl-13 NLS]* | SE170 | SE150 | KB184 | / |
| SGR85 | *greEx71[sur-5p::Hu smad-4 NES::PCKRS; rpr-* | SE170 | SE290 | KB184 | / |

| | | | | | |
|---|---|---|---|---|---|
| | *1p::tRNA(M15/M7)*$_{UCUA}$*; rps-0p::GFP(TAGA)::mCherry::HA::egl-13 NLS]* | | | | |
| SGR86 | *greEx72[sur-5p::Hu smad-4 NES::PCKRS; rpr-1p::tRNA(M15/M7)*$_{UCUA}$*; rps-0p::GFP(TAGA)::mCherry::HA::egl-13 NLS]* | SE170 | SE290 | KB184 | / |
| SGR87 | *greEx73[sur-5p::Hu smad-4 NES::PCKRS; rpr-1p::tRNA(C15/M7)*$_{UCUA}$*; rps-0p::GFP(TAGA)::mCherry::HA::egl-13 NLS]* | SE170 | SE294 | KB184 | / |
| SGR88 | *greEx74[sur-5p::Hu smad-4 NES::PCKRS; rpr-1p::tRNA(C15/M7)*$_{UCUA}$*; rps-0p::GFP(TAGA)::mCherry::HA::egl-13 NLS]* | SE170 | SE294 | KB184 | / |
| SGR89 | *greEx75[glr-1p::Hu smad-4 NES::PCKRS; rpr-1p::tRNA(M15/M7)*$_{UCUA}$*; glr-1p::Cre K201TAGA::SL2::GFP; glr-1p::loxP::B-gal terminator + loxP::ChR2::mKate2]* | SE174 | SE290 | SE368 | IR361 |
| SGR90 | *greEx76[glr-1p::Hu smad-4 NES::PCKRS; rpr-1p::tRNA(M15/UAGA-1)*$_{UCUA}$*; glr-1p::Cre K201TAGA::SL2::GFP; glr-1p::loxP::B-gal terminator + loxP::ChR2::mKate2]* | SE174 | SE302 | SE368 | IR361 |
| SGR91 | *greEx77[sur-5p::Hu smad-4 NES::PCCRS; rpr-1p::tRNA(M15)*$_{CUA}$*; rps-0p::GFP(TAG)::mCherry::HA::egl-13 NLS]* | SE367 | SE149 | SG88 | / |
| SGR92 | *greEx78[sur-5p::Hu smad-4 NES::PCCRS; rpr-1p::tRNA(M15)*$_{CUA}$*; rps-0p::GFP(TAG)::mCherry::HA::egl-13 NLS]* | SE367 | SE149 | SG88 | / |
| SGR93 | *greEx79[sur-5p::Hu smad-4 NES::PCCRS; rpr-1p::tRNA(M15/M7)*$_{UCUA}$*; rps-0p::GFP(TAGA)::mCherry::HA::egl-13 NLS]* | SE367 | SE290 | KB184 | / |
| SGR94 | *greEx80[sur-5p::Hu smad-4 NES::PCCRS; rpr-1p::tRNA(M15/M7)*$_{UCUA}$*; rps-0p::GFP(TAGA)::mCherry::HA::egl-13 NLS]* | SE367 | SE290 | KB184 | / |
| SGR95 | *greEx81[mec-4p::Hu smad-4 NES::PCCRS::SL2::CFP; rpr-1p::tRNA(M15/M7)*$_{UCUA}$*; mec-4p::rev-Caspase-3(CeOpt) C271TAGA]* | ZX334 | SE290 | ZX360 | / |

### Supplementary Table 3.1 - Expression plasmids

| Name | Description | Notes |
|---|---|---|
| IR361 | *glr-1p::loxP::B-gal 3'UTR::loxP::ChR2-mKate2::let-858* | [2] |

| | | |
|---|---|---|
| | *3'UTR* | |
| KB184 | *rps-0p::GFP(TAGA)::mCherry::HA::egl-13 NLS::unc-54 3'UTR* | SG576 + EH5 + SE187; pDEST R4-R3 II |
| SE170 | *sur-5p::hu smad4 NES::Mm PCKRS CeOpt::let-858 3'UTR* | [2] |
| SE174 | *glr-1p::hu smad4 NES::Mm PCKRS CeOpt::let-858 3' UTR* | [2] |
| SE367 | *sur-5p::hu smad4 NES::Mm PCCRS CeOpt::let-858 3'UTR* | SE72 + KB124 + SG606; IR98 |
| SE368 | *glr-1p::PC-Cre K201TAGA::SL2::GFP::let-858 3'UTR* | IR157 + LD441 + IR182; pDEST R4-R3 II |
| SG88 | *rps-0p::GFP(TAG)::mCherry::HA::egl-13 NLS::unc-54 3'UTR* | [3] |
| ZX334 | *mec-4p::hu smad4 NES::Mm PCCRS CeOpt::SL2::GFP::let858 3'UTR* | LD400 + KB124 + ZX114; pDEST R4-R3 II |
| ZX360 | *mec-4p::rev-Casp-3(CeOpt) C135TAGA::unc54 3'UTR* | LD400 + ZX339 + ZX358; SE82 |

### Supplementary Table 3.2 - Destination Vectors

| Name | Description | Notes |
|---|---|---|
| IR98 | *pDEST rps-0p::HygR::unc-54 3'UTR* | [4] |
| SE82 | *pDEST wars-1p::HygR::unc-54 3'UTR* | *rps-0*p was replaced by *wars-1*p amplified from genomic DNA with primers *wars-1*p attB1R (GGGGACTGCTTTTTTGTACAAACTTGGCTGTGTTGAACCCTGAAAAAAATAAATTGGGG) & *wars-1*p attB4F (GGGGACAACTTTGTATAGAAAAGTTGactagtAAGAGCCACCACCGAAATAGATG) |

### Supplementary Table 3.3 - pENTR P4-P1r Vectors

| Name | Description | Notes |
|---|---|---|
| IR157 | *glr-1p* | [2] |
| LD400 | *mec-4p* | Cloned from genomic DNA with primers attB4 *mec-4*p F (GGGGACAACTTTGTATAGAAAAGTTGAACTGCCAATCTGTGCAAATTCAGG) & attB1r *mec-4*p R (GGGGACTGCTTTTTTGTACAAACTTGTCTATAACTTGATAGCGATAAAAAAAATAGCATTAGCAAACTG) |
| SE72 | *sur-5p* | [2] |
| SG576 | *rps-0p* | [5] |

### Supplementary Table 3.4 - pENTR 221 Vectors

| Name | Description | Notes |
|---|---|---|
| EH5 | *GFP-TAGA* | Made from SG4 [3] using primers 457 (ACGAGCTCTACAAGGGAGGATTTaAACCCAGCTTTCTTGTACAA) & 462 (ACGAGCTCTACAAGGGAGGACTCaAACCCAGCTTTCTTGTACAA) |
| KB124 | *hu smad-4 NES::PCCRS Mm CeOpt* | Made from a synthetic gene containing the PCC mutations and a N-terminals Smad4 NES (GCCTGCCCAGTCCCACTCCAACTCCCACCACTCGAGCGTCTCACCCTC GAC) |
| LD441 | *PC-Cre(K201TAGA)* | Made from SG617 [2] by inserting an A to the 5' of amber codon TAG |

| | | |
|---|---|---|
| SE154 | *hu smad4 NES::Mm PCKRS CeOpt* | [2] |
| SG322 | *rpr-1p::PylT:: sup-7 3'* | [2] |
| ZX339 | *rev-Caspase-3(C271TAGA)* | Synthesised after *C. elegans* optimisation, inserted into pDONR221 |

### Supplementary Table 3.5 - pENTR P2r-P3 Vectors

| Name | Description | Notes |
|---|---|---|
| IR182 | *SL2::GFP::let-858 3'UTR* | [2] |
| SE149 | *rpr-1p::tRNA(M15)$_{CUA}$::sup-7 3'* | The expression cassette from SG322 was cloned into a P2r-P3 vector and PylT was replaced with tRNA(M15)$_{CUA}$ (sequence: GGAAACCTGGTCAGGGAGACCGAACGGACTCTAAATCCGTTCAGCCGGGTTCGATTCCCGGGGTTTCCG) |
| SE150 | *rpr-1p::tRNA(C15)$_{CUA}$::sup-7 3'* | The expression cassette from SG322 was cloned into a P2r-P3 vector and PylT was replaced with tRNA(C15)$_{CUA}$ (sequence: GGGAGAGTGGCCAAGGTGGCCGTGTTGACTCTAAATCAACACAGGGGGGTTCGATTCCCCCCTCTCCCG) [2] |
| SE187 | *mCherry CeOpt::HA::egl-13 NLS::unc-54 3'UTR* | Sequence of mCherry was optimised for *C. elegans*, HA and NLS from SG88 was fused to the mCherry 3' end by overlap extension PCR |
| SE290 | *rpr-1p::tRNA(M15/M7)$_{UCUA}$::sup-7 3'* | Made from SE149 using PCR followed by NEBuilder, using primers 602 (CGAAGGGGCTtCTAATCCGCTTCAGCCGGGTTcGATTCC) & 603 (GCGGATTAGaAGCCCCTTCGgTCTcCcTGAcCAGG) |
| SE292 | *rpr-1p::tRNA(M15)$_{UCUA}$::sup-7 3'* | Made from SE149 using PCR followed by NEBuilder, using primers 606 (AAcGGACTtctaAATCCGTTCAGCCGGGTTcG) & 607 (AACGGATTtagaAGTCCgTTCGgTCTcCcTG) |
| SE294 | *rpr-1p::tRNA(C15/M7)$_{UCUA}$::sup-7 3'* | Made from SE150 using PCR followed by NEBuilder, using primers 610 (TGGGGGCTTCTAATCCGCCACAGGGGGGTTCGATTCC) & 611 (TGGCGGATTAGAAGCCCCCACGGCCACCTTGGCCACTC) |
| SE296 | *rpr-1p::tRNA(C15)$_{UCUA}$::sup-7 3'* | Made from SE150 using PCR followed by NEBuilder, using primers 614 (TGTTGACTtctaAATCAACACAGGGGGGTTC) & 615 (TGTTGATTtagaAGTCAACACGGCCACCTTG) |
| SE301 | *rpr-1p::tRNA(Pyl/UAGA-1/)$_{UCUA}$::sup-7 3'* | The expression cassette from SG322 was cloned into a P2r-P3 vector and the anticodon changed using primers 627 (AATGGGCTtctaATCCTGTTCAGCCGGGTTAGATTCCCGGGGTTTC) & 628 (GAACAGGATtagaAGCCCATTCGATCTACATGATC) |
| SE302 | *rpr-1p::tRNA(M15/UAGA-1)$_{UCUA}$::sup-7 3'* | Made from SE149 using PCR followed by NEBuilder, using primers 629 (AAcGGGCTtctaATCCTGTTCAGCCGGGTTcGATTCCCGGGGTTTC) & 630 (AACAGGATtagaAGCCCgTTCGgTCTcCcTGAcCAGGTTTC) |
| SE303 | *rpr-1p::tRNA(C15/UAGA-1)$_{UCUA}$::sup-7 3'* | Made from SE150 using PCR followed by NEBuilder, using primers 631 (CGTGTGGGCTtctaATCCTACACAGGGGGGTTCGATTC) & 632 (TGTAGGATtagaAGCCCACACGGCCACCTTGGCCACTC) |
| SE304 | *rpr-1p::tRNA(Pyl)$_{UCUA}$::sup-7 3'* | The expression cassette from SG322 was cloned into a P2r-P3 vector with anticodon loop changed, using primers 625 (AATGGACTtCTAAATCCGTTCAGCCGGGTTAG) & 626 (GAACGGATTTAGaAGTCCATTCGATCTACATGATC) |
| SG304 | *GFP::let-858 3'UTR* | [2] |
| SG606 | *let-858 3'UTR* | [2] |
| ZX114 | *SL2::CFP::let858 3'UTR* | The *gpd-2/3* intergenic region SL2 was amplified from genomic DNA and fused to optimised CFP. A *let-858* 3' UTR from SG606 was attached to the 3' end of CFP. |

| | | |
|---|---|---|
| ZX358 | *unc-54 3′UTR* | Inserted into pDONR P2R-P3 |

**Supplementary Table 4 - Gene sequences**

The codons used to designate the ncAA incorporation site are marked in red. Different DNA elements are shaded in different colours as indicated. Artificial introns are in lowercase.

| Name & Description | Sequence |
|---|---|
| *Mm PCCRS CeOpt.* (*Methanosarcina mazei* pyrrolysine aminoacyl tRNA synthetase mutated to recognise photocaged cysteine and codon optimised for expression in *C. elegans*) | ATGGACAAGAAGCCACTCAACACCCTCATCTCCGCCACCGGACTCT<br>GGATGTCCCGTACCGGAACCATCCACAAGATCAAGCACCACGAGGT<br>CTCCCGTTCCAAGATCTACATCGAGATGGCCTGCGGAGACCACCTC<br>GTCGTCAACAACTCCCGTTCCTCCCGTACCGCCCGTGCCCTCCGTC<br>ACCACAAGTACCGTAAGACCTGCAAGCGTTGCCGTGTCTCCGACGA<br>GGACCTCAACAAGTTCCTCACCAAGGCCAACGAGGACCAAACCTCC<br>GTCAAGGTCAAGGTCGTCTCCGCCCCAACCCGTACCAAGAAGgtaa<br>gtttaaacatatatatactaactaaccctgattatttaaattttca<br>gGCCATGCCAAAGTCCGTCGCCCGTGCCCCAAAGCCACTCGAGAAC<br>ACCGAGGCCGCCCAAGCCCAACCATCCGGATCCAAGTTCTCCCCAG<br>CCATCCCAGTCTCCACCCAAGAGTCCGTCTCCGTCCCAGCCTCCGT<br>CTCCACCTCCATCTCCTCCATCTCCACCGGAGCCACCGCCTCCGCC<br>CTCGTCAAGGGAAACACCAACCCAATCACCTCCATGTCCGCCCCAG<br>TCCAAGCCTCCGCCCCAGCCCTCACCAAGTCCCAAACCGACCGTCT<br>CGAGGTCCTCCTCAACCCAAAGGACGAGATCTCCCTCAACTCCGGA<br>AAGCCATTCCGTGAGCTCGAGTCCGAGCTCCTCTCCCGTCGTAAGg<br>taagtttaaacagttcggtactaactaaccatacatatttaaattt<br>tcagAAGGACCTCCAACAAATCTACGCCGAGGAGCGTGAGAACTAC<br>CTCGGAAAGCTCGAGCGTGAGATCACCCGTTTCTTCGTCGACCGTG<br>GATTCCTCGAGATCAAGTCCCCAATCCTCATCCCACTCGAGTACAT<br>CGAGCGTATGGGAATCGACAACGACACCGAGCTCTCCAAGCAAATC<br>TTCCGTGTCGACAAGAACTTCTGCCTCCGTCCAATGCTCGCCCCAA<br>ACCTCTACAACTACCTCCGTAAGCTCGACCGTGCCCTCCCAGACCC<br>AATCAAGATCTTCGAGATCGGACCATGCTACCGTAAGGAGTCCGAC<br>GGAAAGgtaagtttaaacatgattttactaactaactaatctgatt<br>taaattttcagGAGCACCTCGAGGAGTTCACCATGCTCCAGTTCGC<br>CCAAATGGGATCCGGATGCACCCGTGAGAACCTCGAGTCCATCATC<br>ACCGACTTCCTCAACCACCTCGGAATCGACTTCAAGATCGTCGGAG<br>ACTCCTGCATGGTCTACGGAGACACCCTCGACGTCATGCACGGAGA<br>CCTCGAGCTCTCCTCCGCCATGGTCGGACCAATCCCACTCGACCGT<br>GAGTGGGGAATCGACAAGCCATGGATCGGAGCCGGATTCGGACTCG<br>AGCGTCTCCTCAAGGTCAAGCACGACTTCAAGAACATCAAGCGTGC<br>CGCCCGTTCCGAGTCCTACTACAACGGAATCTCCACCAACCTCTAA |
| *Mm PCKRS CeOpt*. (*Methanosarcina mazei* pyrrolysine aminoacyl tRNA synthetase mutated to recognise photocaged lysine and codon optimised for expression in *C. elegans*) | ATGGACAAGAAGCCACTCAACACCCTCATCTCCGCCACCGGACTCT<br>GGATGTCCCGTACCGGAACCATCCACAAGATCAAGCACCACGAGGT<br>CTCCCGTTCCAAGATCTACATCGAGATGGCCTGCGGAGACCACCTC<br>GTCGTCAACAACTCCCGTTCCTCCCGTACCGCCCGTGCCCTCCGTC<br>ACCACAAGTACCGTAAGACCTGCAAGCGTTGCCGTGTCTCCGACGA<br>GGACCTCAACAAGTTCCTCACCAAGGCCAACGAGGACCAAACCTCC<br>GTCAAGGTCAAGGTCGTCTCCGCCCCAACCCGTACCAAGAAGgtaa<br>gtttaaacatatatatactaactaaccctgattatttaaattttca<br>gGCCATGCCAAAGTCCGTCGCCCGTGCCCCAAAGCCACTCGAGAAC<br>ACCGAGGCCGCCCAAGCCCAACCATCCGGATCCAAGTTCTCCCCAG<br>CCATCCCAGTCTCCACCCAAGAGTCCGTCTCCGTCCCAGCCTCCGT<br>CTCCACCTCCATCTCCTCCATCTCCACCGGAGCCACCGCCTCCGCC<br>CTCGTCAAGGGAAACACCAACCCAATCACCTCCATGTCCGCCCCAG<br>TCCAAGCCTCCGCCCCAGCCCTCACCAAGTCCCAAACCGACCGTCT<br>CGAGGTCCTCCTCAACCCAAAGGACGAGATCTCCCTCAACTCCGGA<br>AAGCCATTCCGTGAGCTCGAGTCCGAGCTCCTCTCCCGTCGTAAGg<br>taagtttaaacagttcggtactaactaaccatacatatttaaattt<br>tcagAAGGACCTCCAACAAATCTACGCCGAGGAGCGTGAGAACTAC |

| | |
|---|---|
| | CTCGGAAAGCTCGAGCGTGAGATCACCCGTTTCTTCGTCGACCGTG<br>GATTCCTCGAGATCAAGTCCCCAATCCTCATCCCACTCGAGTACAT<br>CGAGCGTTTCGGAATCGACAACGACACCGAGCTCTCCAAGCAAATC<br>TTCCGTGTCGACAAGAACTTCTGCCTCCGTCCAATGCTCTCCCCAA<br>ACCTCTGCAACTACATGCGTAAGCTCGACCGTGCCCTCCCAGACCC<br>AATCAAGATCTTCGAGATCGGACCATGCTACCGTAAGGAGTCCGAC<br>GGAAAGgtaagtttaaacatgattttactaactaactaatctgatt<br>taaattttcagGAGCACCTCGAGGAGTTCACCATGCTCAACTTCTG<br>CCAAATGGGATCCGGATGCACCCGTGAGAACCTCGAGTCCATCATC<br>ACCGACTTCCTCAACCACCTCGGAATCGACTTCAAGATCGTCGGAG<br>ACTCCTGCATGGTCTACGGAGACACCCTCGACGTCATGCACGGAGA<br>CCTCGAGCTCTCCTCCGCCGTCGTCGGACCAATCCCACTCGACCGT<br>GAGTGGGGAATCGACAAGCCATGGATCGGAGCCGGATTCGGACTCG<br>AGCGTCTCCTCAAGGTCAAGCACGACTTCAAGAACATCAAGCGTGC<br>CGCCCGTTCCGAGTCCTACTACAACGGAATCTCCACCAACCTCTAA |
| *GFP(TAG)::mCherry::HA::egl-13 NLS*. (GFP, **amber stop codon**, mCherry, HA, and NLS of *egl-13* linked in tandem) | ATGTCCAAGGGAGAGGAGCTCTTCACCGGAGTCGTCCCAATCCTCG<br>TCGAGCTCGACGGAGACGTCAACGGACACAAGTTCTCCGTCTCCGG<br>AGAGGGAGAGGGAGACGCCACCTACGGAAAGCTCACCCTCAAGTTC<br>ATCTGCACCACCGGAAAGCTCCCAGTCCCATGGCCAACCCTCGTCA<br>CCACCTTCACCTACGGAGTCCAATGCTTCTCCCGTTACCCAGgtaa<br>gtttaaacatatatatactaactaaccctgattatttaaattttca<br>gACCACATGAAGCGTCACGACTTCTTCAAGTCCGCCATGCCAGAGG<br>GATACGTCCAAGAGCGTACCATCTTCTTCAAGGACGACGGAAACTA<br>CAAGACCCGTGCCGAGGTCAAGTTCGAGGGAGACACCCTCGTCAAC<br>CGTATCGAGCTCAAGGGAATCGACTTCAAGgtaagtttaaacagtt<br>cggtactaactaaccatacatatttaaattttcagGAGGACGGAAA<br>CATCCTCGGACACAAGCTCGAGTACAACTACAACTCCCACAACGTC<br>TACATCATGGCCGACAAGCAAAAGAACGGAATCAAGGTCAACTTCA<br>AGATCCGTCACAACATCGAGGACGGATCTGTCCAACTCGCCGACCA<br>CTACCAACAAAACACCCCAATCGGAGACGGACCAGTCCTCCTCCCA<br>Ggtaagtttaaacatgattttactaactaactaatctgatttaaat<br>tttcagACAACCACTACCTCTCCACCCAATCCGCCCTCTCCAAGGA<br>CCCAAACGAGAAGCGTGACCACATGGTCCTCAAGGAGTTCGTCACC<br>GCCGCCGGAATCACCCACGGAATGGACGAGCTCTACAAGGGAGGA**T**<br>**AG**GGCGCGCCAGGCCGGCCAAACCCAGCTTTCTTGTACAAAGTGGC<br>CATGGTCTCAAAGGGTGAAGAAGATAACATGGCAATTATTAAAGAG<br>TTTATGCGTTTCAAGGTGCATATGGAGGGATCTGTCAATGGGCATG<br>AGTTTGAAATTGAAGGTGAAGGAGAAGGCCGACCATATGAGGGAAC<br>ACAAACCGCAAAACTAAAGgtaagtttaaacatatatatactaact<br>aaccctgattatttaaattttcagGTAACTAAAGGCGGACCATTAC<br>CATTCGCCTGGGACATCCTCTCTCCACAGTTCATGTATGGAAGTAA<br>AGCTTATGTTAAACATCCGGCAGATATACCAGATTATTTGAAACTT<br>TCATTCCCGGAGGGTTTTAAGTGGGAACGCGTAATGAATTTTGAAG<br>ACGGAGGAGTTGTTACAGTGACGCAAGACTCAAGgtaagtttaaac<br>agttcggtactaactaaccatacatatttaaattttcagCCTCCAA<br>GATGGAGAATTTATTTATAAAGTCAAACTTCGAGGAACGAATTTCC<br>CCTCGGATGGACCTGTTATGCAGAAGAAGACTATGGGATGGGAAGC<br>TTCAAGTGAAAGAATGTACCCTGAAGACGGTGCTCTTAAGGGAGAG<br>ATTAAACAACGTCTTAAATTGAAAGATGGAGGACATTACGATGCTG<br>AGgtaagtttaaacatgattttactaactaactaatctgatttaaa<br>ttttcagGTGAAGACAACTTACAAAGCCAAAAAACCAGTTCAGCTG<br>CCAGGAGCGTACAATGTTAATATTAAACTGGATATCACCTCCCACA<br>ACGAGGATTACACTATCGTTGAGCAATATGAAAGAGCTGAAGGGCG<br>GCACTCGACAGGTGGCATGGATGAATTGTATAAGTACCCATATGAT<br>GTCCCAGACTACGCTATGAGCCGTAGACGAAAAGCGAATCCGACAA<br>AACTGAGTGAAAACGCGAAGAAGCTTGCCAAGGAAGTTGAAAATTA<br>A |
| *GFP(TAGA)::mCherry::HA::egl-13 NLS*. (GFP, **quadruplet codon TAGA**, mCherry, HA, and NLS of *egl-13* linked in tandem) | ATGTCCAAGGGAGAGGAGCTCTTCACCGGAGTCGTCCCAATCCTCG<br>TCGAGCTCGACGGAGACGTCAACGGACACAAGTTCTCCGTCTCCGG<br>AGAGGGAGAGGGAGACGCCACCTACGGAAAGCTCACCCTCAAGTTC<br>ATCTGCACCACCGGAAAGCTCCCAGTCCCATGGCCAACCCTCGTCA<br>CCACCTTCACCTACGGAGTCCAATGCTTCTCCCGTTACCCAGgtaa<br>gtttaaacatatatatactaactaaccctgattatttaaattttca |

| | |
|---|---|
| | gACCACATGAAGCGTCACGACTTCTTCAAGTCCGCCATGCCAGAGG<br>GATACGTCCAAGAGCGTACCATCTTCTTCAAGGACGACGGAAACTA<br>CAAGACCCGTGCCGAGGTCAAGTTCGAGGGAGACACCCTCGTCAAC<br>CGTATCGAGCTCAAGGGAATCGACTTCAAGGTAAGTTTAAACAGTT<br>CGGTACTAACTAACCATACATATTTAAATTTTCAGGAGGACGGAAA<br>CATCCTCGGACACAAGCTCGAGTACAACTACAACTCCCACAACGTC<br>TACATCATGGCCGACAAGCAAAAGAACGGAATCAAGGTCAACTTCA<br>AGATCCGTCACAACATCGAGGACGGATCTGTCCAACTCGCCGACCA<br>CTACCAACAAAACACCCCAATCGGAGACGGACCAGTCCTCCTCCCA<br>Ggtaagtttaaacatgattttactaactaactaatctgatttaaat<br>tttcagACAACCACTACCTCTCCACCCAATCCGCCCTCTCCAAGGA<br>CCCAAACGAGAAGCGTGACCACATGGTCCTCAAGGAGTTCGTCACC<br>GCCGCCGGAATCACCCACGGAATGGACGAGCTCTACAAGGGAGGA**T**<br>**AGA**AACCCAGCTTTCTTGTACAAAGTGGGAATGGTCTCCAAGGGAG<br>AGGAGGACAACATGGCCATCATCAAGGAGTTCATGCGTTTCAAGGT<br>CCACATGGAGGGATCCGTCAACGGACACGAGTTCGAGATCGAGGGA<br>GAGGGAGAGGGACGTCCATACGAGGGAACCCAAACCGCCAAGCTCA<br>AGGTCACCAAGGTAAGTTTAATCATATATATACTAACTAACCCTGA<br>TTATTTAAATTTTCAGGGAGGACCACTCCCATTCGCCTGGGACATC<br>CTCTCCCCACAATTCATGTACGGATCCAAGGCCTACGTCAAGCACC<br>CAGCCGACATCCCAGACTACCTCAAGCTCTCCTTCCCAGAGGGATT<br>CAAGTGGGAGCGTGTCATGAACTTCGAGGACGGAGGAGTCGTCACC<br>GTCACCCAAGACTCCTCCCTCCAAGACGGAGAGTTCATCTACAAGG<br>TAAGTTTAAACAGTTCGGTACTAACTAACCATACATATTTAAATTT<br>TCAGGTCAAGCTCCGTGGAACCAACTTCCCATCCGACGGACCAGTC<br>ATGCAAAAGAAGACCATGGGATGGGAGGCCTCCTCCGAGCGTATGT<br>ACCCAGAGGACGGAGCCCTCAAGGGAGAGATCAAGCAACGTCTCAA<br>GCTCAAGgtaagtttaaacatgattttactaactaactaatctgat<br>ttaaattttcagGACGGAGGACACTACGACGCCGAGGTCAAGACCA<br>CCTACAAGGCCAAGAAGCCAGTCCAACTCCCAGGAGCCTACAACGT<br>CAACATCAAGCTCGACATCACCTCCCACAACGAGGACTACACCATC<br>GTCGAGCAATACGAGCGTGCCGAGGGACGTCACTCCACCGGAGGAA<br>TGGACGAGCTCTACAAGTACCCATACGACGTCCCAGACTACGCCAT<br>GTCCCGTCGTCGAAAGGCCAACCCAACCAAGCTCTCCGAGAACGCC<br>AAGAAGCTCGCCAAGGAGGTCGAGAAC |
| *PC-Cre(K201TAG)*. (*C. elegans* codon optimised Cre recombinase with K201 replaced by an **amber stop codon**) | ATGGGGGCGCCAAAAAAGAAGAGGAAAGTATCGAATTTGCTCACCG<br>TCCACCAAAACCTCCCAGCCCTCCCAGTCGACGCCACCTCCGACGA<br>GGTCCGTAAGAACCTCATGGACATGTTCCGTGACCGTCAAGCCTTC<br>TCCGAGCACACCTGGAAGATGCTCCTCTCCGTCTGCCGTTCCTGGG<br>CCGCCTGGTGCAAGCTCAACAACCGTAAGTGGTTCCCAGCCGAGCC<br>AGAGGACGTCCGTGACTACCTCCTCTACCTCCAAGCCCGTGGACTC<br>GCCGTCAAGgtaagtttaaacatatatatactaactaaccctgatt<br>atttaaattttcagACCATCCAACAACACCTCGGACAACTCAACAT<br>GCTCCACCGTCGTTCCGGACTCCCACGTCCATCCGACTCCAACGCC<br>GTCTCCCTCGTCATGCGTCGTATCCGTAAGGAGAACGTCGACGCCG<br>GAGAGCGTGCCAAGCAAGCCCTCGCCTTCGAGCGTACCGACTTCGA<br>CCAAGTCCGTTCCCTCATGGAGAACTCCGACCGTTGCCAAGACATC<br>CGTAACCTCGCCTTCCTCGGAATCGCCTACAACACCCTCCTCCGTA<br>TCGCCGAGATCGCCCGTATCCGTGTCAAGGTAAGTTTAAACAGTTC<br>GGTACTAACTAACCATACATATTTAAATTTTCAGGACATCTCCCGT<br>ACCGACGGAGGACGTATGCTCATCCACATCGGACGTACC**TAG**ACCC<br>TCGTCTCCACCGCCGGAGTCGAGAAGGCCCTCTCCCTCGGAGTCAC<br>CAAGCTCGTCGAGCGTTGGATCTCCGTCTCCGGAGTCGCCGACGAC<br>CCAAACAACTACCTCTTCTGCCGTGTCCGTAAGgtaagtttaaaca<br>tgattttactaactaactaatctgatttaaattttcagAACGGAGT<br>CGCCGCCCCATCCGCCACCTCCCAACTCTCCACCCGTGCCCTCGAG<br>GGAATCTTCGAGGCCACCCACCGTCTCATCTACGGAGCCAAGGACG<br>ACTCCGGACAACGTTACCTCGCCTGGTCCGGACACTCCGCCCGTGT<br>CGGAGCCGCCCGTGACATGGCCCGTGCCGGAGTCTCCATCCCAGAG<br>ATCATGCAAGCCGGAGGATGGACCAACGTCAACATCGTCATGAACT<br>ACATCCGTAACCTCGACTCCGAGACCGGAGCCATGGTCCGTCTCCT<br>CGAGGACGGAGACTAA |

| | |
|---|---|
| *PC-Cre(K201TAGA)*. (*C. elegans* codon optimised Cre recombinase with K201 replaced by a **quadruplet codon TAGA**) | ATGGGGGCGCCAAAAAAGAAGAGGAAAGTATCGAATTTGCTCACCG<br>TCCACCAAAACCTCCCAGCCCTCCCAGTCGACGCCACCTCCGACGA<br>GGTCCGTAAGAACCTCATGGACATGTTCCGTGACCGTCAAGCCTTC<br>TCCGAGCACACCTGGAAGATGCTCCTCTCCGTCTGCCGTTCCTGGG<br>CCGCCTGGTGCAAGCTCAACAACCGTAAGTGGTTCCCAGCCGAGCC<br>AGAGGACGTCCGTGACTACCTCCTCTACCTCCAAGCCCGTGGACTC<br>GCCGTCAAGgtaagtttaaacatatatatactaactaaccctgatt<br>atttaaattttcagACCATCCAACAACACCTCGGACAACTCAACAT<br>GCTCCACCGTCGTTCCGGACTCCCACGTCCATCCGACTCCAACGCC<br>GTCTCCCTCGTCATGCGTCGTATCCGTAAGGAGAACGTCGACGCCG<br>GAGAGCGTGCCAAGCAAGCCCTCGCCTTCGAGCGTACCGACTTCGA<br>CCAAGTCCGTTCCCTCATGGAGAACTCCGACCGTTGCCAAGACATC<br>CGTAACCTCGCCTTCCTCGGAATCGCCTACAACACCCTCCTCCGTA<br>TCGCCGAGATCGCCCGTATCCGTGTCAAGgtaagtttaaacagttc<br>ggtactaactaaccatacatatttaaattttcagGACATCTCCCGT<br>ACCGACGGAGGACGTATGCTCATCCACATCGGACGTACC**TAGA**ACC<br>CTCGTCTCCACCGCCGGAGTCGAGAAGGCCCTCTCCCTCGGAGTCA<br>CCAAGCTCGTCGAGCGTTGGATCTCCGTCTCCGGAGTCGCCGACGA<br>CCCAAACAACTACCTCTTCTGCCGTGTCCGTAAGgtaagtttaaac<br>atgattttactaactaactaatctgatttaaattttcagAACGGAG<br>TCGCCGCCCCATCCGCCACCTCCCAACTCTCCACCCGTGCCCTCGA<br>GGGAATCTTCGAGGCCACCCACCGTCTCATCTACGGGGCCAAGGAC<br>GACTCCGGACAACGTTACCTCGCCTGGTCCGGACACTCCGCCCGTG<br>TCGGAGCCGCCCGTGACATGGCCCGTGCCGGAGTCTCCATCCCAGA<br>GATCATGCAAGCCGGAGGATGGACCAACGTCAACATCGTCATGAAC<br>TACATCCGTAACCTCGACTCCGAGACCGGAGCCATGGTCCGTCTCC<br>TCGAGGACGGAGACTAA |
| *rev-Caspase-3(C271TAGA)*. (*C. elegans* codon optimised subunit-reversed Caspase-3 with C271 replaced by a **quadruplet codon TAGA**) | ATGAGCGGGGTGGATGATGATATGGCTTGTCACAAGATCCCAGTCG<br>AGGCCGACTTCCTCTACGCCTACTCCACCGCCCCAGGATACTACTC<br>CTGGCGTAACTCCAAGGACGGATCCTGGTTCATCCAATCCCTCTGC<br>GCCATGCTCAAGCAATACGCCGACAAGCTCGAGTTCATGCACATCC<br>TCACCCGTGTCAACCGTAAGGTCGCCACCGAGTTCGAGTCCTTCTC<br>CTTCGACGCCACCTTCCACGCCAAGGTAAGTTTAAACATATCTATA<br>CTAACTAACCCTGATTATTTAAATTTTCAGAAGCAAATCCCATGCA<br>TCGTCTCCATGCTCACCAAGGAGCTCTACTTCTACCACGACGAGGT<br>CGACGGAATGGAGAACACCGAGAACTCCGTCGACTCCAAGTCCATC<br>AAGAACCTCGAGCCAAAGATCATCCACGGATCCGAGTCCATGGACT<br>CCGGAATCTCCCTCGACAACTCCTACAAGATGGACTACCCAGAGAT<br>GGGACTCTGCATCATCATCAACAACAAGgtaagtttaaacagttcg<br>gtactaactaaccatacatatttaaattttcagAACTTCCACAAGT<br>CCACCGGAATGACCTCCCGTTCCGGAACCGACGTCGACGCCGCCAA<br>CCTCCGTGAGACCTTCCGTAACCTCAAGTACGAGGTCCGTAACAAG<br>AACGACCTCACCCGTGAGGAGATCGTCGAGCTCATGCGTGACGTCT<br>CCAAGGAGGACCACTCCAAGCGTTCCTCCTTCGTCTGCGTCCTCCT<br>CTCCCACGGAGAGGAGGGAATCATCTTCGGAACCAACGGACCAGTC<br>GACCTCAAGgtaagtttaaacatgattttactaactaactaatctg<br>atttaaattttcagAAGATCACCAACTTCTTCCGTGGAGACCGTTG<br>CCGTTCCCTCACCGGAAAGCCAAAGCTCTTCATCATCCAAGCC**TAG**<br>**A**CGTGGAACCGAGCTCGACTGCGGAATCGAGACCGACTAA |


1. Srinivasula, S.M., et al., *Generation of constitutively active recombinant caspases-3 and -6 by rearrangement of their subunits.* J Biol Chem 1998. **273**(17): p. 10107-11.
2. Davis, L., et al., *Precise optical control of gene expression in C. elegans using genetic code expansion and Cre recombinase*, U.o. Edinburgh, Editor. 2020: http://www.bioRxiv.org.

3. Greiss, S. and J.W. Chin, *Expanding the genetic code of an animal.* J Am Chem Soc, 2011. **133**(36): p. 14196-9.
4. Radman, I., S. Greiss, and J.W. Chin, *Efficient and rapid C. elegans transgenesis by bombardment and hygromycin B selection.* PLoS One, 2013. **8**(10): p. e76019.
5. Hunt-Newbury, R., et al., *High-Throughput In Vivo Analysis of Gene Expression in Caenorhabditis elegans.* PLoS Biology, 2007. **5**(9): p. e237.
